# Plasticity of extrachromosomal and intrachromosomal BRAF amplifications in mediating targeted therapy dosage challenges

**DOI:** 10.1101/2021.11.23.468420

**Authors:** Kai Song, Jenna K. Minami, William P. Crosson, Jesus Salazar, Eli Pazol, Niroshi T. Senaratne, Nagesh Rao, Kim Paraiso, Thomas G. Graeber

## Abstract

Cancer cells display two modes of focal amplifications (FAs), extrachromosomal DNA/double-minutes (ecDNA/DMs) and intrachromosomal homogenously staining regions (HSRs). Understanding the plasticity of these two modes is critical for preventing targeted therapy resistance. We developed a combined BRAF plus MEK inhibitor resistance melanoma model that bears high BRAF amplifications through both DM and HSR modes, and investigated FA dynamics in the context of drug resistance plasticity. Cells harboring FAs displayed mode switching between DMs and HSRs, from both de novo genetic changes and selection of preexisting subpopulations. We found that copy number plasticity is not exclusive to DMs. Single cell-derived clones with HSRs also exhibit BRAF copy number and corresponding HSR length plasticity that allows them to respond to dose reduction and recover from drug addiction. Upon kinase inhibitor escalation, we observed reproducible selection for cells with BRAF kinase domain duplications residing on DMs. In sum, the plasticity of FAs allows cancer cells to respond to drug dose changes through a myriad of mechanisms. These mechanisms include increases or decreases in DMs, shortening of HSRs, acquisition of secondary resistance mechanisms, and expression of alternative slicing oncogene variants. These results highlight the challenges in targeting the cellular vulnerabilities tied to focal amplifications.

**Statement of Significance:** Understanding the dynamics of oncogene amplifications is critical for appreciating tumorigenesis and preventing anticancer drug resistance. We found melanoma cells harboring BRAF amplifications in either DM or HSR formats to have high plasticity under different kinase inhibitor dosage challenges with evidence supporting de novo alterations, clonal selection, and coupling to additional resistance mechanisms. In in the absence of DMs, HSRs can offer comparable levels of plasticity as DMs.

## Introduction

Genomic instability, an important enabling characteristic in cancer, confers cells with a list of aberrant hallmarks such as enhanced invasion and deregulated cellular energetics ^1^. Among many types of instability-driven mutations, focal amplifications (FAs) of oncogenes in cancer genomes is a major contributor of neoplastic progression and therapeutic resistance ^2–5^. There are primarily two modes of FAs: double minute (DM) and homogeneously staining region (HSR). DMs are circular extrachromosomal DNAs (ecDNAs) which allow copies of oncogenes to exist freely in nuclei and retain intact and even elevated transcription activity ^6–9^. DMs are able to replicate autonomously, but are acentric and therefore segregate into daughter cells randomly ^10^. HSRs are intrachromosomal amplifications caused by tandem duplications of oncogene-containing regions resulting in long segments with uniform staining intensities in cytogenetics ^11^. Several models regarding the generation of these two kinds of FAs have been proposed, including but not restricted to breakage-fusion-bridge, episomal and chromothripsis mechanisms ^12–15^. DMs can be an intermediary state leading to HSRs via chromosomal re-integration ^16,17^. The high prevalence of both kinds of FAs support their importance in tumorigenesis ^18^. DMs have been observed in large number of tumors of different types, especially in neuroblastomas (31.0%) and adrenal tumors (27.6%), but rarely in normal tissues. A high occurrence of the HSR form of FAs is found in particular cancer types such as squamous cell carcinoma (12.1%) and oral cavity (10.9%), but across all cancers HSRs have a slightly lower frequency compared to DMs (0.8% vs 1.3%, respectively) ^19^. FAs are present more frequently in complex neoplastic systems such as patient tumor and PDX, rather than cell lines ^18^.

Mutations in BRAF, a serine/threonine RAF family kinase and a key upstream member of the MAPK pathway, have been associated with many cancer types. The frequency of BRAF mutations varies widely across cancer types. For example, BRAF mutations are relatively common in thyroid gland and skin cancers (60% and 52% respectively), but are very rare in kidney cancers (0.3%) ^20^. In melanoma therapy, the development of BRAF inhibitors, such as vemurafenib and dabrafenib as well as combinatorial treatments with other MAPK pathway inhibitors (e.g. MEK) have greatly improved patient survival ^21^. However, acquired resistance often compromises the efficacy of these therapies. To date, many resistance mechanisms to BRAF inhibition emerging during clinical treatment have been identified, including reactivation of the MAPK pathway, activation of the PI3K/AKT pathway, or both. This can occur via genomic mutations, genomic rearrangements such as kinase domain duplication, altered splice isoform variant expression, cellular dedifferentiation, and other mechanisms ^5,22–29^. One mechanism of reactivating the MAPK pathway that is frequently found in melanoma patient tumors is the acquisition of BRAF amplifications ^5^. As previously noted, these amplifications can be found on DMs and HSRs ^25,30,31^. However, the structure, plasticity and dynamics of BRAF DMs and HSRs as well as the different conditions required for their generation during acquired drug resistance remain unclear, and as such is the focus of our current study.

In this study, through acquired BRAF and MEK inhibitor resistance, we developed a melanoma model system that dynamically harbors BRAF in the form of DMs, HSRs or both. These amplicons were then characterized and revealed that genes adjacent to BRAF were upregulated at both genomic and transcriptomic levels. By establishing and experimenting with single-cell-derived clones, we found that increasing and/or decreasing kinase inhibitor dose is a reproducible method to modulate the number of DMs or the length of HSRs, as well as to influence the transition between these two amplification modes. In our experiments, the resistance-driven population possessed a high degree of heterogeneity in terms of karyotypes, and cells tended to transition from DMs to HSRs when drug dose was held at steady state. Moreover, drug dose challenges generated a subpopulation of cells with BRAF kinase domain duplications (KDD), an additional resistance mechanism for BRAF inhibition. This combined FA karyotype and KDD genotype led to high dose kinase inhibitor tolerance, with reduced DM copy number requirement. Collectively, our findings demonstrate extensive genomic and phenotypic plasticity enabled by or coupled to FAs via DM copy numbers, HSR lengths, transitions between the two modes, and enabling of additional genomic rearrangements.

## Results

### Acquired resistance to BRAF and MEK kinase inhibitors resulted in both DM and HSR karyotypes

In order to generate a FA-positive melanoma model, we treated a human melanoma cell line, M249 with the clinically used kinase inhibitor combination: vemurafenib (BRAF inhibitor, BRAFi) and selumetinib (MEK inhibitor, MEKi) to develop resistance (abbreviated as M249-VSR for <u>v</u>emurafenib and <u>s</u>elumetinib <u>r</u>esistant) as previously described ^32^. The doses for both drugs were sequentially increased by roughly 2-fold, with each dose escalation taking place when cells resumed growth rates with doubling in 4 days or less. The initial and final does were 0.05μM and 2μM, respectively (Fig. 1A). Upon the establishment of cells resistant to 2μM doses, fluorescent in situ hybridization (FISH) was performed and showed a high amplification of BRAF, primarily in the DM form. However, over the course of a few months in culture, these cells spontaneously switched their karyotypes to DM-negative HSR-positive without any observed exceptions (Fig. 1B; see Supplementary Fig. S1 and Methods section for categories and images of BRAF FA FISH-based karyotypes). To quantify the extent of FA, we performed quantitative real time PCR (qPCR) on M249-VSR-DM and -HSR cells and found that there were 30-fold to 40-fold increase in BRAF copy numbers, respectively, compared to M249 parental cells which contained 5 copies of BRAF (Fig.1B and C). Such amplifications also led to high gene expression levels of BRAF (Fig. 1D).

**Figure 1.**
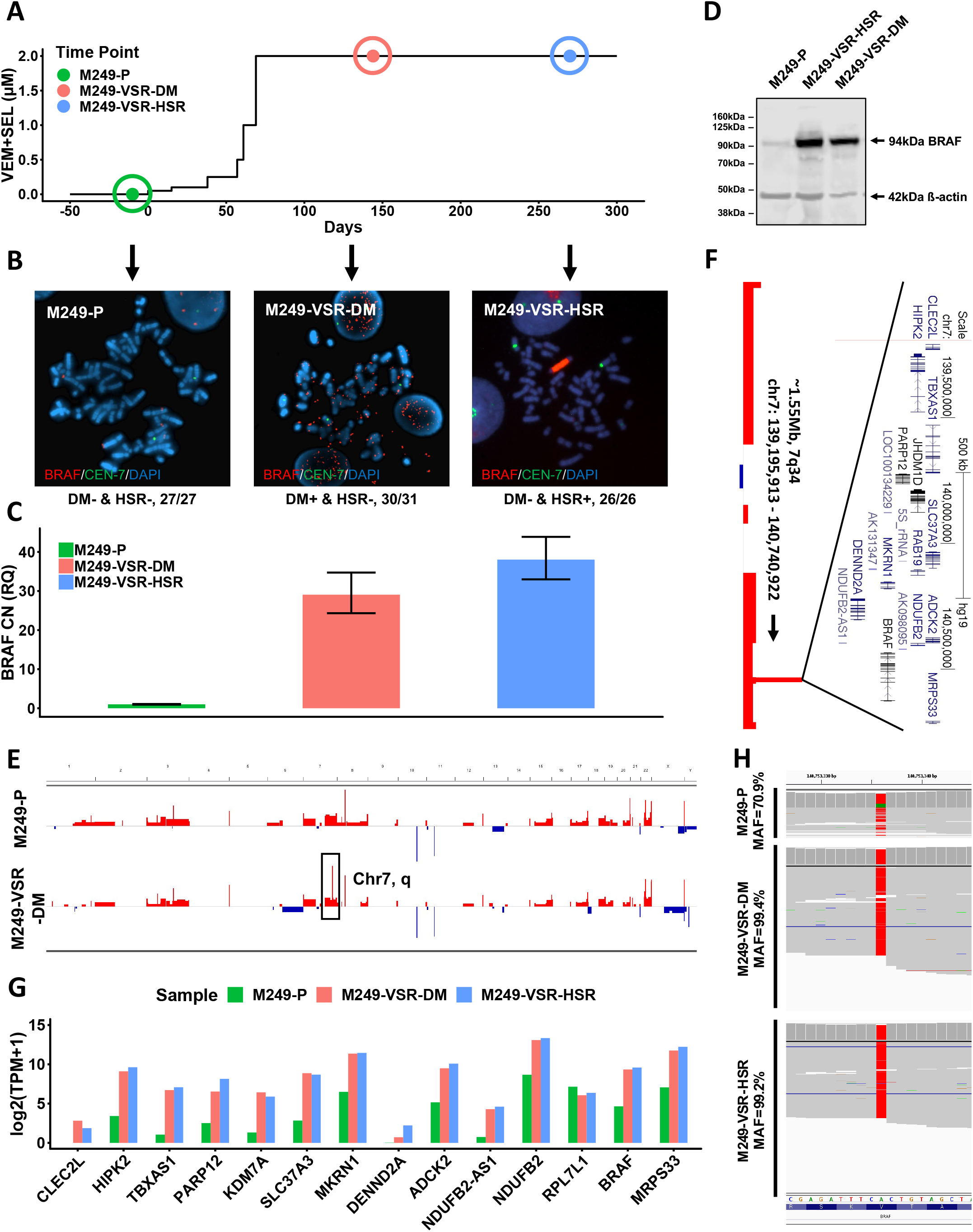
Focal amplifications in the form of DMs and HSRs mediate resistance to BRAF +MEK inhibition. **A**, BRAFi+MEKi treatment history for M249 cells. Circles around the time points represent the rough windows of sample collections (genomic DNA, total RNA, whole cell lysates and fixations for FISH). **B**, FISH images showed three different karyotypes coming from corresponding time points in (**A**). Representative images are only shown if the karyotypes occurred with high percentages. The fraction under each image represents the number of observations for this karyotype divided by total number of observations. CEN-7: centromere 7. **C**, qPCR results of relative BRAF copy number in the samples from three time points in (**A**). All values were normalized to GAPDH copy number of corresponding samples and then to M249 parental cells. Error bars represent t-distribution-based 95% confidence intervals from technical triplicates. CN: copy number. RQ: relative quantity. Three technical replicates. **D**, Immunoblot of BRAF for all three corresponding samples in (**A**). **E**, CGH results of M249-P and M249-VSR-DM showed that the most significant copy number increase happened on 7q34. **F**, Zoom-in of the focal amplified region in (**E**). Gene annotations within the amplicon were obtained from UCSC genome browser. **G**, RNA-seq analysis of the three samples in (**A**) shows that focal amplifications propagated to transcriptomic level. Only genes residing on amplicons were plotted. **H**, Frequencies of c.1799T>A (V600E) in M249-P, -VSR-DM and -VSR-HSR cells. It is inferred by aligning RNA-seq reads to the genome. MAF: major allele frequency. Green: thymine. Red: adenine. See also Supplementary Fig. S2.

Since qPCR is limited to investigating a small DNA region, we employed comparative genomic hybridization (CGH) and whole genome sequencing (WGS) for M249-P and M249-VSR-DM cells to reveal the full copy number alterations (CNA) across the genome (Fig. 1E and Supplementary Fig. S2). Though there were other alterations, the most striking change upon acquisition of resistance was a FA of size ~1.55Mb at chr7q34, the region of the BRAF locus, with a fold increase consistent with qPCR results (log2 ratio = 5.144). Genes adjacent to BRAF on the amplicon were amplified to a similar degree (Fig. 1F); and as measured by RNA-seq, the transcripts of these genes were also elevated (Fig. 1G). Such co-amplifications have also been found on amplicons containing other oncogenes, e.g. MYC and EGFR ^33,34^. A recent study reported that co-amplifying EGFR enhancers with EGFR can contribute to cancer cell fitness ^34^. RNA-seq based SNV calling of DM and HSR M249 cell lines indicated that the BRAF 1799T>A (V600E) mutation, the target of vemurafenib, was selected during FA development, with both DM and HSR cases displaying greater than 99% major allele frequency compared to 71% in the parental line (Fig. 1H). Thus, the focal amplicons in both the DM and HSR versions of M249-VSR contain similar genomic material, a result suggesting reintegration of DMs followed by a selective advantage as the mechanism of transition to the HSR case.

### Single-cell-derived clones reveal de novo integrations of DMs into chromosomes as HSRs

To further dissect changes that occurred during the transition from DMs to HSRs, we isolated five single-cell-derived clones (SCs) from the bulk M249-VSR population. These clones were isolated at an intermediary timepoint, between the DM+ & HSR- and DM- & HSR+ karyotypes (Fig. 2A). Cultures derived from these single cell clones were expanded and characterized for subsequent changes over a three month timeframe. At the outset, three of the resultant clones had a DM+ & HSR- karyotype, one clone had a DM- & HSR+ karyotype, and one had a DM+ & HSR+ karyotype (Fig. 2B and C, SC1-SC5; Supplementary Table S1). Over the three-month time course, the bulk population began with a small percentage of DM- & HSR+ cells which gradually expanded to dominate the population (Fig. 2B and 2C, B1-B4). In contrast, the five SCs displayed a range of evolutionary trajectories. The majority of SC1 cells kept their DM+ & HSR- karyotype, but 4.1% (1/24) of the cells had their DMs integrated as HSRs, and 12.5% (3/24) cells harbored both DMs and HSRs (Fig. 2B, SC1-2). SC4, another line that began as DM+ & HSR-, also developed small portions of DM- & HSR+ (11.1%, 2/18) and DM+ & HSR+ (16.7%, 3/18) karyotypes. Interestingly, HSRs of some SC4 cells presented in a format that had three smaller HSR segments in different chromosomes in each cell. In contrast, in all other cells, one long HSR, with or without an additional short HSR segment, was observed (Fig. 2B, SC4-2). The data above supports that de novo integration of DMs as HSRs did occur. However, no changes occurred in one DM+ & HSR- clone, SC3, indicating that the tendency for integration is not absolute on the time scale observed (Fig. 2B, SC3-2).

**Figure 2.**
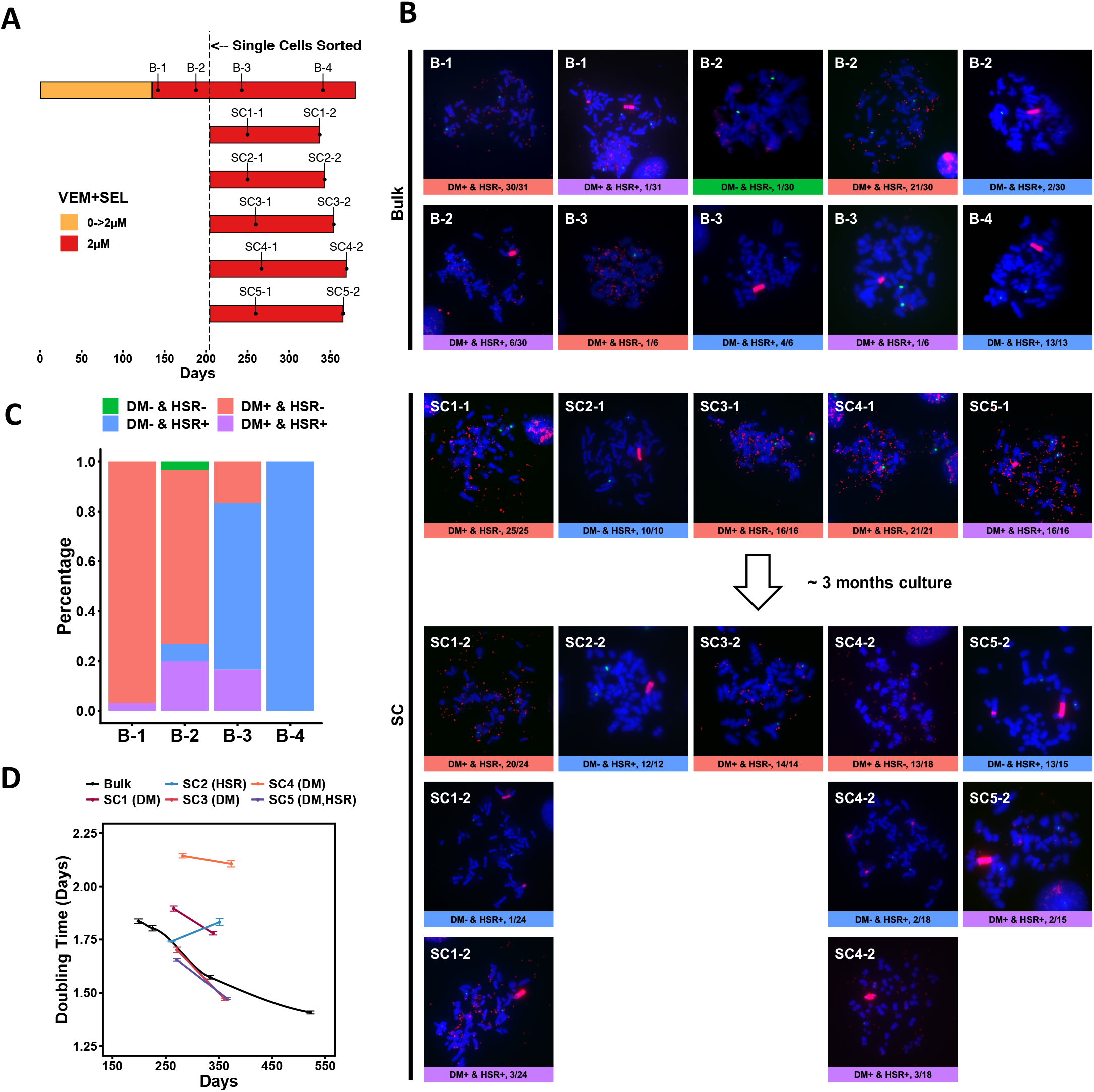
Single-cell-derived clones reveal de novo integrations of DMs into chromosomes as HSRs. **A**, The timeline of deriving M249-VSR single-cell-derived clones (SCs). **B**, FISH images of all sampling points for both bulk and SC samples in (**A**). Red: BRAF. Green: centromere 7. Blue: DAPI. **C**, Plotted are karyotype percentages for all sampling points of bulk M249-VSR cells in (**A**). **D**, Doubling times of all SCs near the sampling points. Sampling points of bulk cells were shown by days. e.g. 199D: 199 days from the start of resistance development of M249 cells, not to be confused with bulk sampling points in (**A**). Error bars represents standard error of means (SEMs) of doubling times (see Methods for SEM calculation). Three technical replicates. See also Supplementary Fig. S3 and S4.

Several findings support the notion that HSRs are a more stable focal amplification mode ^35^. First, clone SC2, which contained only HSRs at the outset, maintained its karyotype and BRAF copy number after three months in culture (Fig. 2B, SC2-2; Supplementary Fig. S3). Second, the most drastic FA mode switch was observed in SC5: 86.7% (13/15) of the cells switched from DM+ & HSR+ to DM- & HSR+, with only 13.3% retaining the mixed karyotype (Figure 2B, SC5-2). As changes occurred rapidly in SC5, cells with DM+ & HSR+ karyotypes appear to reflect an intermediate transition stage in the karyotype switch. These findings are based on single cell clones, and thus directly demonstrate the plasticity of FAs, with a general trend of evolution from DMs to HSRs in this system.

In addition to de novo karyotype changes, we observed growth rate differences before and after the three-month expansion among the single-cell-derived clones. Four out of five SCs, including three DM only (SC1, SC3, SC4) and one DM plus HSR (SC5) clones, displayed continuously increased proliferation rates (decreased doubling times) over the three-month culture. These results generally matched the increased proliferation rate seen in the bulk population. In contrast, the HSR only clone (SC2) did not increase its proliferation rate, but rather demonstrated a decrease during this period (Fig. 2D and Supplementary Fig. S4). The single cell clone studies also suggest that additional modes of adaption can happen in isolated cases. For example, clone SC3 had a substantial reduction in the copy number level of BRAF (Supplementary Fig. S3) but nonetheless increased its proliferation rate. This result suggests that this clone had additional genetic events occur that reduced its reliance on high BRAF copy number.

### Non-steady dose challenge can prolong or prevent DM integration into chromosomes

The observation that in the M249-VSR system DMs will integrate into chromosomes as HSRs upon continuous culture at a constant drug dose, suggests that DM+ cells have a fitness disadvantage compared to HSR+ cells in these conditions. However, DMs are often observed in tumor samples, and thus may have a fitness advantage in other conditions. To test this hypothesis, we thought to investigate a scenario in which DMs could have a fitness advantage. DMs are known to segregate asymmetrically during cell division ^18,36–38^, so we tested whether an oscillating drug dose would give DM+ cells increased fitness, arguably through increased heterogeneity of the population (Fig. 3A, EXP2). We designed an experiment to turn double-drug doses on and off in a cycle of 8 days. DMs were indeed retained at high levels without a switch to an HSR state for a longer period of time compared to the steady dose scenario. In the steady dose case, all cells (26/26) were HSR positive on day 262, but in the oscillating dose case there was no detected HSR positive cells (0/15) even approximately 7 weeks later on day 308 (Fig. 3A-3C, FIX5 and Supplementary Fig. S5). However, the number of DMs did decrease in these cells, and the BRAF copy number declined notably (Fig. 3D). We thus investigated whether another known MAPK inhibitor resistance mechanism had emerged in these cells. We found that these cells express the shorter BRAF splice isoform associated with acquired resistance (Supplementary Fig. S5C) ^24^. Thus, in response to the altered fitness challenge of a regularly changing environment, the emerging cells retained DMs longer than cells experiencing constant drug dose, and also converted their BRAF isoform to one that required a lower amount of expression, and thus lower DM numbers.

**Figure 3.**
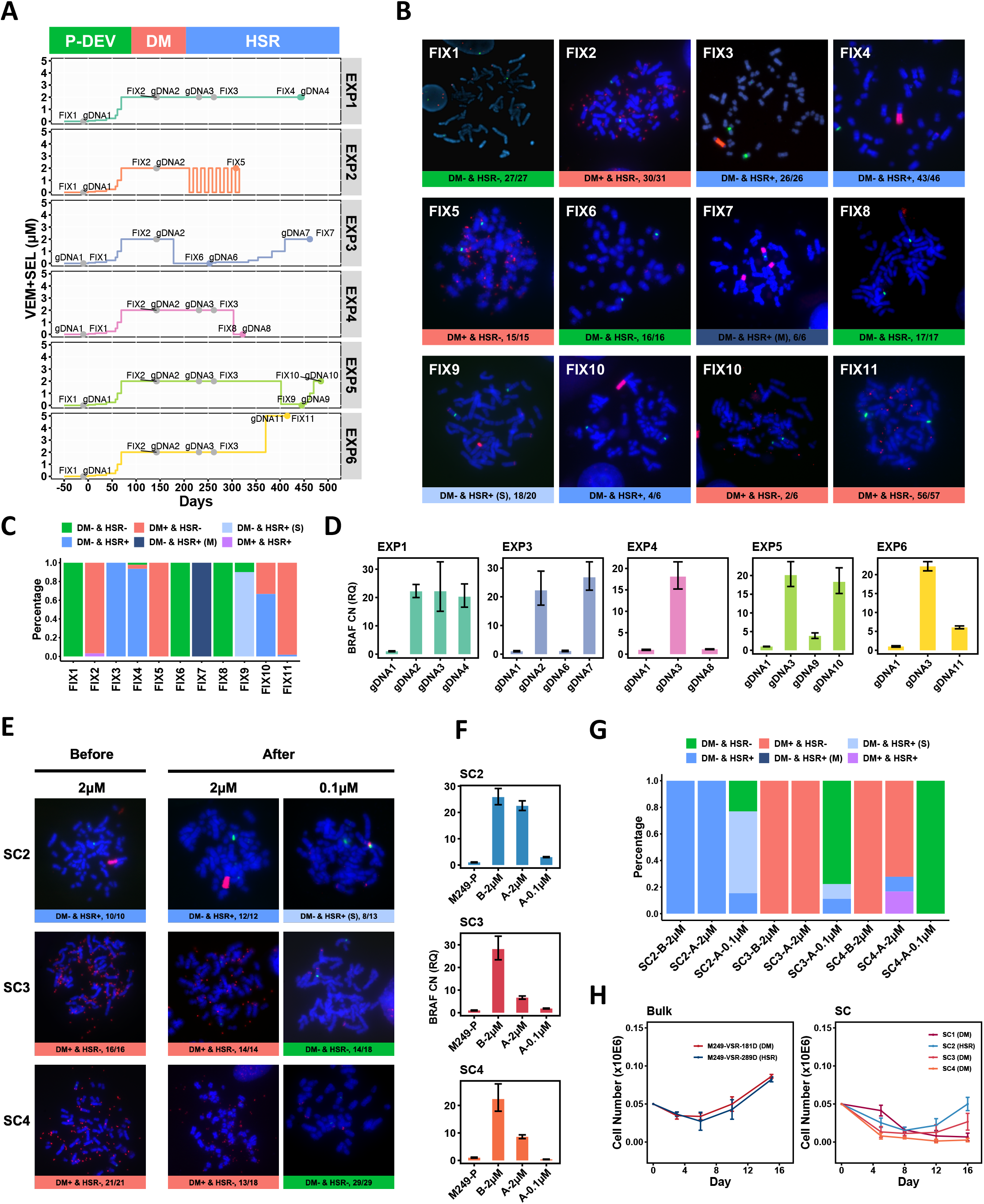
A variety of focal amplifications modes and secondary resistance mechanisms mediate dynamic plasticity to BRAF +MEK inhibition. **A**, The double-drug treatment schemes of various experiments for bulk cells with labels of time points for when cells were fixed (FIX) for FISH and their genomic DNA (gDNA) were extracted. Top bar shows the estimated duration of each stage, inferred from (Fig. 1B) and (Fig. 2B). P-DEV: resistance development from M249 parental cells. DM: the stage when the karyotype is predominantly DM+ & HSR-. HSR: the stage when the karyotype is predominantly DM- & HSR+. Grey dots represent some common time points between different experiments. **B**, Representative FISH images of all fixation points in (**A**). Images are only shown if the corresponding karyotypes occurred at high percentages. Red: BRAF. Green: centromere 7. Blue: DAPI. **C**, Full karyotype percentages of samples in (**A**). DM- & HSR+ (M): multiple HSRs; DM- & HSR+ (S): short HSRs. **D**, Representative qPCR results of BRAF copy number for some gDNA extraction points in (**A**). All values were normalized to GAPDH copy number of corresponding samples and then to gDNA1 (M249 parental cells). Error bars represent t-distribution-based 95% confidence intervals from technical triplicates. **E,** Representative FISH images of three single-cell-derived clones that were treated either with 2μM (original dose) or 0.1μM VEM+SEL for roughly three months. Red: BRAF. Green: centromere 7. Blue: DAPI. **F**, qPCR of the samples in (**E**), all values were normalized to GAPDH copy number of corresponding samples and then to M249-P (M249 parental cells). Error bars represent t-distribution-based 95% confidence intervals from three technical triplicates. B: before. A: after three-month culture. **G**, The full percentage of each karyotype in each sample in (**E**). **H**, Graphed are cell numbers after VEM+SEL was withdrawn from M249-VSR bulk cells and single-cell-derived clones. Error bars are standard deviations from three technical replicates. Predominant BRAF FA modes are denoted in parenthesis. See also Supplementary Fig. S5-S7.

### Both BRAF DM and HSR containing cells display dynamic plasticity upon changes in drug dose

Next, we focused on studying the plasticity of DMs and HSRs in M249-VSR cells. As a foundation for this analysis, we first examined whether HSRs were the final stable form of amplicons for cells kept under constant drug dose by checking their karyotypes after a few additional months. We found that most cells still harbored HSRs with similar amplicon length and BRAF copy number (Fig. 3A-3D, EXP1, sample 3 vs sample 4). This stable result provides our reference control for comparison to other cases with drug dose manipulation.

To evaluate if the DM to HSR trajectory observed under constant inhibitor dose could be affected by changes in dosing, we next either decreased or elevated the double-drug concentration being applied to DM+ or HSR+ cells. Previous studies have examined the potential of using drug removal to eliminate drug-addicted cells ^28,31,32,39^, thus sparking our interest in studying the effect of this approach on DMs and HSRs.

To investigate this, we withdrew VEM+SEL treatment from M249-VSR-DM and -HSR cells. In the DM+ case, when doses were acutely brought down from 2μM to 0μM, all DMs were eliminated based on FISH analysis with the fastest change observed in 12 days. qPCR results showed that the copy number of BRAF was reduced many folds compared to parental BRAF copy number (Fig. 3A-3D, EXP3; Supplementary Fig. S6). The majority of the cells contained DMs prior to drug removal. It is possible that these rapid changes were due to selection of a preexisting DM-negative subpopulation. Other possible explanations include that DMs were exported out of cells through micronuclei exclusions, or that post-mitosis cells with less DMs due to uneven segregation during division could have been selected for upon drug withdrawal ^18,36–38,40^. A prompt reversion of BRAF copy number to the parental state also occurred in HSR cells upon double-drug removal (Fig. 3A-3D, EXP4). Notably, there was not a substantial difference between the recovery time of DM and HSR cells in these drug wash-off experiments.

We repeated the dose decrease experiment above using the bulk population in its HSR+ state. But compared to the experiment described above, the drug reduction was not a complete withdrawal (Fig. 3A-3D, EXP5: 2μM to 0.1μM). In this experiment, the bulk population demonstrated a substantial shortening of the typical HSR length, but HSRs were still detectable. Using this new sub-population, we further explored the cellular genomic plasticity by subsequently reinstating the 2μM double drug dose. The cells display resistance in less than a month, and most cells again presented with the longer form of HSRs (Fig. 3A-3C, EXP5). During the interval of drug reduction and increase, BRAF DNA copy number also decreased following the 2μM to 0.1μM transition, and re-increased following the 0.1μM to 2μM transition accordingly (Fig. 3D, EXP5).

We also elevated the reinstatement of a 2μM drug dose on the bulk population cells that had drug withdrawal (0μM) occur while they were in the DM+ state (EXP3). In this case, it took about 4 months for the cells to re-develop resistance to VEM+SEL (Fig. 3A, EXP3), similar to the time required for the initial establishment of resistance in the parental cells. In this experiment, the melanoma cells demonstrated an additional variation in that upon becoming resistant they typically harbored three separate shorter HSRs on different chromosomes (Fig. 3B and 3C, EXP3). None of the cells presented with either DMs or with a single larger HSR. This treatment course thus further revealed the plasticity of genomic options available for adjusting to changes in selection pressures.

The dose decrease and increase results above could be explained by selection for residual BRAF copy number-low or -high cells in the respective populations. To investigate cellular plasticity to dramatic drug reduction in a more homogeneous population, we turned to the single cell-derived DM+ or HSR+ clones. In these experiments we lowered the VEM+SEL double dose from 2μM to 0.1μM using clones SC2 (HSR), SC3 (DM) and SC4 (DM). In the post-drug-decrease populations, SC2 showed reduced length of HSRs, and SC3 and SC4 showed reduced number of DMs. All cases were accompanied by a substantial drop of BRAF copy number (Fig. 3E-3G). Another characteristic that indicates the plasticity of HSR-harboring cells is comparable to that of the DM case is that the recovery times upon dose withdrawal for DM+ or HSR+ cells, either bulk or as SCs, were not substantially different (Fig. 3H).

These single cell clone results suggest that de novo genetic alterations occur during expansion form a single cell, and/or during the stress of drug withdrawal, thus creating population heterogeneity and enabling population plasticity. In these cases, selection alone cannot explain the outcome, and clearly genomic instability is diversifying the population. DM+ cells would be expected to diversify with every cell division. The result is less anticipated for HSR+ single cell clones due to the focal amplification existing on a chromosome. HSR+ cells must have distinct mechanisms from DM+ cells for re-generating heterogeneity either after single cell cloning or during the stress of drug withdrawal.

While the bulk and single cell clone experiments above demonstrate the plasticity of the HSR form of FAs, we identified other cases of cell lines with HSRs that were not readily tunable by changing BRAFi+MEKi doses. The cell line M395-VSR-HSR was also both i) initially derived by our protocol of elevating drug dose over time, and ii) its resistant state was associated with the presence of a BRAF locus containing HSR. However, when we removed drug from M395-VSR-HSR, the HSRs did not substantially shorten or disappear (Supplementary Fig. S7).

### Karyotypic shift from HSRs to DMs carrying BRAF kinase domain duplications upon double-drug dose increase

In an additional experiment, we further tested the plasticity of melanoma cells harboring focal amplifications. In this case, we elevated the double MAPK inhibitor doses applied to the bulk M249-VSR cells at a timepoint when they were predominantly HSR+. This treatment converted the population of predominantly HSR+ cells to predominantly DM+ cells (Fig. 3A-3C, EXP6). Contrary to the expectation that in the higher drug dose the cells would have higher levels of BRAF DNA copy number, we found that the copy number had decreased approximately 4-fold based on a genomic DNA qPCR assay (Fig. 3D).

To investigate this change further, we repeated the experiment using bulk M249-VSR-HSR cells at various time points over the entire HSR-harboring period, roughly 260 days onwards from the beginning of resistance development (Fig. 4A). Four out of five dose-increase experiments resulted in changes of FA types from HSRs to DMs (VS5-1, VS5-2, VS5-3, VS5-4 and VS5-6 (5 sampling points in total)). One out of five resulted in cells that were DM- and HSR- VS5-5) (Fig. 4B and 4C). Upon further investigation, we found that the five DM+ 5μM-resistant samples all expressed a BRAF protein variant with a molecular weight of approximately 140kDa (Fig. 4D). Four of the five DM+ 5μM-resistant samples also expressed the 62kD variant of BRAF, the BRAF inhibitor-resistant splice variant observed in the oscillating dose experiment above. The 140kD size matches a previously reported BRAF variant with a kinase domain duplication (KDD) that leads to BRAF inhibitor resistance ^25,41,42^. To assay for KDD in our samples, we used a RT-qPCR primer pair that spans the BRAF exon 18-10 junction as described by Kemper et al., 2016. We discovered that all of the HSR to DM transformed samples carried exon 18-10 junctions, while other cultures, including M249 parental (VS0), M249-HSR cells prior to dose increase (VS2-1, VS2-2, VS2-3 and VS2-4) and M249-HSR cells that showed DM and HSR double negative post dose increase (VS5-5), contained none or only a minimal amount of such junctions (Fig. 4E and 4F).

**Figure 4.**
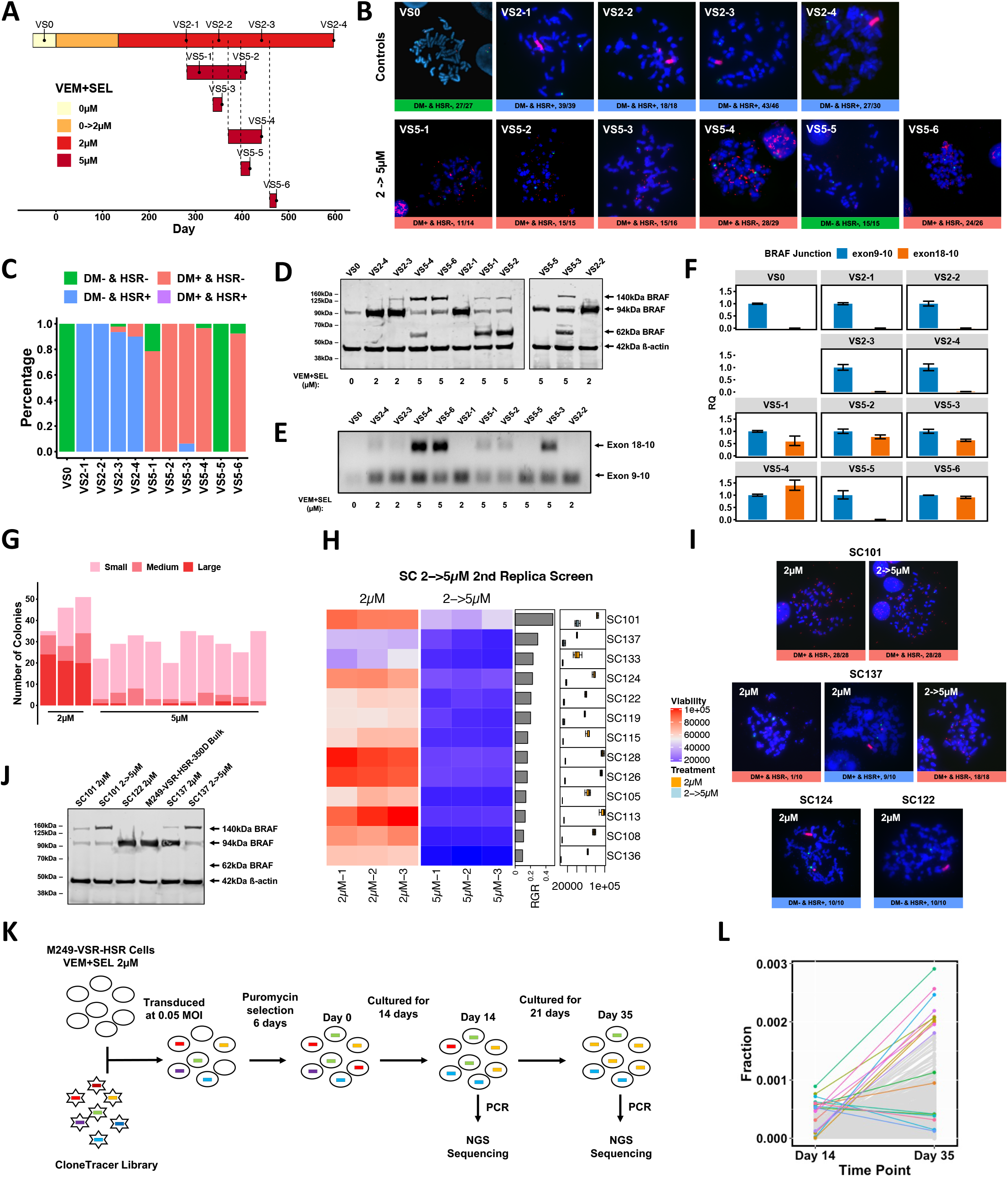
HSR to DM karyotypic switching and BRAF kinase domain duplications mediate resistance to MAPK inhibitor dose increase. **A**, The relationship between samples examined during the processes of M249 VSR development and VEM+SEL 2μM to 5μM dose increases. Cells were collected when they were able to double in 4 days under 5μM dose. **B**, Representative FISH images of all samples in (**A**). Red: BRAF. Green: centromere 7. Blue: DAPI. **C**, The summary of all karyotypes in each sample in (**A**). **D**, Immunoblot of all samples in (**A**), using an antibody that targets the N-terminus of BRAF (12-156aa). The 130kD band is the KDD form, and the 62kD band is the alternatively spliced form of BRAF. **E and F**, qPCR and RT-PCR for all samples in (**A**) with primer sets that target BRAF exon 18-10 and exon 9-10 junctions. For RT-qPCR, all values of exon junction 18-10 were normalized to that of exon junction 9-10 of corresponding samples. Error bars represent SEMs around ΔCt values derived by Satterthwaite approximation. **G**, M249-VSR-HSR bulk cells were sorted into single cells (on day 322 using the scale in Fig. 3A) and seeded in 13 96-well plates. Next they were treated with either original 2μM (n=3) or 5μM VEM+SEL (n=10) for 12 days. The sizes of the resultant colonies were classified into three categories (Small, Medium and Large) by eye. **H**, second replica screen for M249-VSR-HSR single-cell-derived clones that tolerate VEM+SEL 2 to 5μM dose increase. Rows of the heatmap represent different clones ranked by relative growth rate (RGR), calculated by dividing the mean at 5μM by that at 2μM. **I**, representative FISH images of selected clones in (**H**) with estimated percentage of each kind of FA. Red: BRAF. Green: centromere 7. Blue: DAPI. **J**, Immunoblot of BRAF in bulk cells and singlecell-derived clones treated with the indicated dose regiments. **K**, Design of the barcode-based clone tracing experiments. Cells were transduced with the lentivirus ClonTracer library on day 318 based on the timeline in (**A**). **L**, Comparison between barcode fractions on Day 14 and Day 35 as depicted in (**K**). Top 10 barcodes by fraction from each sampling time point are highlighted. See also Supplementary Fig. S8-S10.

We next investigated whether the KDD was developed due to selection of an existing subpopulation or de novo kinase domain duplications only after 2 to 5μM dose increase. Under constant 2μM dose VEM+SEL, the M249-VSR resistant cells were initially primarily DM+ & HSR- (circa day 150), turned primarily DM- & HSR+ with time (circa day 260), and then with additional time reacquired a small percentage of DM+ & HSR- cells (450 days and onwards, Fig. 4A-4C). Their late timepoint DM+ & HSR- fractions were 2/46 (4.35%, VS2-3) and 3/30 (10%, VS2-4) respectively. This expanding DM+ population could have been the source of KDD that expanded post drug dose increase to 5μM.

To further test if rare DM+ cells were present at earlier times below the level of detection by bulk FISH analysis, we used both single cell sorting and a replica plating approach. First, cells from the earlier-stage M249-VSR-HSR bulk population (322 days) were single cell sorted. This collection of single cell clones was then either treated at the original 2μM dose or at an elevated 5μM dose of VEM+SEL (Fig. 4G; Supplementary Fig. S8A). We found that 3.2% of the single cell clones could grow under 5μM in a similar manner compared to their counterparts at 2μM (percentage calculated as the number of large colonies per plate at 5μM divided by the number at 2μM).

Next we added a replica plating step. Forty-one of the single-cell clones derived at 2μM were replica plated, and then treated in parallel at either the original 2μM dose or the elevated 5μM dose (Fig. 4H; schematic of protocol in Supplementary Fig. S8). After two rounds of screening, the fastest growing clone (SC101), as calculated by dividing the growth rate at 5μM by that at 2μM, was revealed to be DM+ & HSR- both before and after the dose increase, with no observed cellular heterogeneity of FA modes (Fig. 4I). The second fastest clone (SC137) started with a 10% (1/10) DM+ & HSR- population, but finished at 100% (18/18) DM+ & HSR- at the end of the 2 rounds of 5μM screening (Fig. 4I). Four other randomly selected SCs from either the near-top of the relative growth rate-sorted list and the bottom of the list displayed no DM+ & HSR- karyotype (SC122, SC124, SC111, SC106, Fig. 4I and Supplementary Fig. S8C). The two fastest SCs SC101 and SC137 did harbor BRAF KDD on their DMs according to immunoblot analysis (Fig. 4J). In a second quantitative viability assay, the SC101 and SC137 KDD+ SCs again demonstrated the best ability to tolerate drug dose increases (Supplementary Fig. S9). This replica screening result supports that cells harboring the BRAF KDD containing DMs preexisted in the bulk population prior to increases in the drug dose. These cells were starting to expand in the 2μM drug condition, but the increase in drug dose sharply increased any relative fitness advantage.

We next designed additional experiments to distinguish between clonal selection versus de novo generation of KDD-bearing DMs upon dose escalation to 5μM. First, we used the single-cell-derived clones to test if dose escalation can readily convert HSRs to KDD-bearing DMs. Longterm treatment of single-cell-derived DM- HSR+ M249-VSR clones SC2 and SC208 with 5μM VEM+SEL resulted in growing cells, but failed to generate a DM+ population (Supplementary Fig. S10A and S10B). Second, we used a barcode-based clone tracing system (ClonTracer) ^43^ to keep track of the subpopulations in the bulk M249-VSR-HSR cells (from day 318). Even under the constant 2μM drug dose, certain cells did expand faster than the rest (Fig. 4K and 4L). This results demonstrates that even at this later timepoint and under steady drug dose exposure, the bulk population of cells continues to evolve in regards to its subpopulation distributions.

## Discussion

Focal amplifications of oncogenes in either DM- (ecDNA-) or HSR-mode are clinically observed both in the treatment-naïve setting (e.g. MYCN in neuroblastoma ^17^), and as a resistance mechanism for inhibitors targeting these oncogenes (e.g. erlotinib treatments in glioblastoma ^44^). The disappearance of oncogene-containing DMs has also been reported upon modeling of oncogene-targeted therapy ^45^. While a few-fold amplification of BRAF is sometimes observed in treatment-naïve melanoma tumors ^46^, higher-fold DM- or HSR-mode focal amplifications are typically seen only following BRAF inhibitor therapy ^30,32^. To further elucidate the genomic plasticity enabled by focal amplifications, we developed an expanded version of a BRAF kinase inhibition and BRAF locus amplification model. This system demonstrated a high degree and broad range of evolutionary plasticity in response to changing drug dose regiments. This plasticity included multiple genomic rearrangement and related mechanisms such as kinase domain duplications and alternative splicing.

In the initial phase of drug resistance to dual BRAF and MEK inhibition, the BRAF amplification appeared via DMs. Under conditions of a stable double drug dose, the population gradually became dominated by an HSR-form of BRAF amplification. The DM to HSR conversion was also observed in single-cell-derived clones of the M249-VSR cells, supporting that integration was a de novo event during evolution, rather than both DMs and HSRs co-occurring in the generation of the resistant subline. This result adds to reports in the literature. For example, one study conducted a long-term observation on a non-drug-treated leukemia cell line and saw formation of MYC-carrying HSRs from DMs ^35^. Other studies have inferred a common origin and mode switch interpretation of data that showed high similarities between DM and HSR subclones of multiple parental cell lines ^47,48^.

The reproducible observation of the DM to HSR transition led us to hypothesize that DMs carry a higher fitness disadvantage than HSRs during stable conditions. In support of this, we found that an oscillating drug dose could prevent or prolong autosomal integration of the amplicon. This difference in fitness is arguably linked to uneven segregation of DMs ^18,38,49^, and the resulting uneven *BRAF* gene copy numbers in daughter cells. During non-stable conditions, cellular heterogeneity provided by uneven segregation can provide a reservoir of cells more adept to grow well in the new conditions. In contrast, during stable drug-dose conditions a reduction in cellular heterogeneity would produce a fitness advantage, which all daughter cells maintaining the optimal *BRAF* gene copy number. We did observe one DM-carrying SC that did not switch to HSR positive during long term stable culture. In this case, the DM copy number of these cells decreased, suggesting that a secondary (undetermined) resistance mechanism allowed these cells to depend less on the heterogeneous DMs (Fig. 2B and Supplementary Fig. S3). Taken together, our results support that in stable contexts DMs are not a fitness optimized form of amplification, and thus tend to be replaced by other mechanisms such as less heterogeneous chromosomally integrated HSRs.

Nevertheless, our study revealed that ‘HSR plasticity’ can be a mode of tumor evolution in response to drug challenge. We are unaware of previous studies that have directly made similar HSR plasticity observations. The uneven segregation of ecDNAs provides an evidence-supported model for tumor heterogeneity and thus tumor ability to withstand changes in targeted drug dosing ^18,38,49^. Our study uncovered that in the absence of DMs, HSRs can offer somewhat comparable levels of plasticity as DMs. Due to the inherent differences between DM and HSR modes of amplification, this is almost undoubtedly through distinct molecular mechanisms. In more detail, we observed single-cell-derived HSR-containing cell populations that demonstrated dose-tunable HSR lengths (Fig. 3E). This plasticity resembles the dose-tunable feature of DM- harboring populations. In the single-cell-derived HSR+ experiments we did not observe any DMs. This argues against a model in which the long HSRs were first excised and shredded from chromosomes to form multiple DMs and then re-integrated back to chromosomes as short HSRs. Future work will investigate whether errors and repairs made while replicating and segregating intrachromosomal long HSRs may be generating heterogeneity and thus contributing to this plasticity.

The tumor evolutionary and drug resistance plasticity enabled by focal amplifications extended beyond changes in amplicon copy numbers and DM versus HSR modes. In particular we observed two additional parallel mechanisms, i) kinase domain duplication, representing an additional genomic rearrangement mechanism ^25,41,42^, and ii) activation of an alternative splicing mechanism ^24^. Our results indicate cells harboring BRAF KDD-encoded DMs and/or the alternative splicing mechanism can be reproducibly selected from an HSR-predominant population upon drug escalation treatment. The data supports an interpretation in which the cells with KDD-harboring DMs remained within the HSR-harboring cell population, and upon drug escalation the KDD-harboring cells gained a relative fitness advantage. Further research on KDD formation and KDD-mediated resistance could offer therapeutic insights for pan-cancer therapy, as this alteration occurs to many other kinases, such as EGFR and FGFR1 in glioma and lung cancer ^50–53^. We also observed the alternative splicing mechanism as a potential method to escape reliance on high DM copy number during an oscillating dose regiment (Fig. 3A, EXP2). The drug resistance provided by the splice variant, arguably lowers the number of DMs required, but maintains the DM-mediated unequal segregation-based heterogeneity.

Therapeutic approaches to target the vulnerabilities of FA-harboring cells, as well as other genomic instability-associated vulnerabilities, are in academic and industry development. Our study demonstrates important challenges, such as mode switching and acquisition of additional genomic rearrangements, that must be co-addressed in these pursuits. Previous reports have revealed that different amplification modes can have associated targetable vulnerabilities, as well ss distinct mechanisms for FA generation and maintenance. For example, treating cells with ribonucleotide reductase inhibitors like hydroxyurea or gemcitabine removes DMs ^54,55^ but not HSRs ^56^. Suppressing homologous recombination by silencing BRCA1 inhibits DM-containing cells, while displaying minimal effects on HSR-containing counterparts ^57^. Furthermore, overexpression of Sei-1 promotes DM formation ^58^; ERK1/2 activation contributes to DM production ^59^; and transducing p53 null mouse ovarian surface epithelial cells with myristoylated-AKT and KRAS G12D construct generates MYC DMs *in vivo* ^60^.

Collectively, we observed a high degree and broad range of tumor evolution and drug resistance plasticity enabled by or coupled to focal amplifications. Through perturbations by a panel of drug regiment challenges, we observed i) de novo generation of extrachromosomal DMs, ii) de novo integration of DMs into chromosomal HSRs, iii) context-dependent HSR-mediated fitness advantage over DMs, iv) context-dependent DM-mediated fitness advantage over HSRs, v) co-evolution of DMs and a de novo genomic rearrangement creating a kinase domain duplication, vi) co-evolution of DMs and activation of BRAF alternative splicing, vii) propensity to couple secondary resistance mechanisms (KDD and/or alternative splicing) to DMs to reduce the total number of DMs required, and viii) a plasticity of HSRs that compares to the known plasticity of DMs and can result in de novo conversion of long HSRs to short HSRs. Appreciation of the interplay of focal amplification modes with drug regiments and other resistance mechanisms is central to our understanding of tumor evolution and drug resistance, and to developing therapeutic approaches to overcome the resulting plasticity.

## Methods

### Cell Culture Conditions and Generation of Drug-resistant Cell Lines

The M249 (RRID: CVCL_D755) and M395 (RRID: CVCL_XJ99) cell lines are part of the M series melanoma lines established from patient biopsies at UCLA under UCLA IRB approval #02-08-06 and were obtained from Antoni Ribas ^61^. All cells were cultured in RPMI 1640 with L-glutamine (Gibco), 10% (v/v) fetal bovine serum (Omega Scientific), and 1% (v/v) streptomycin (Gibco). All cells were maintained in a humidified 5% CO2 incubator. BRAF inhibitor vemurafenib and MEK inhibitor selumetinib were obtained from Selleckchem and their stock solutions were prepared in DMSO. Resistance cell lines were generated by exposing cells to step-wise increasing doses of vemurafenib and selumetinib, similar to the previously described approach ^32^. Growth, inhibition of growth and death were assayed by staining cells with trypan blue (Sigma-Aldrich) followed by cell counting using Vi-cell XR Cell Viability Analyzer (Beckman Coulter).

### Single-cell-derived Clones and Flow Cytometry

Resistant subclones were derived by seeding single cells from the bulk population into 96-well plates using FACSAria cell sorter. Doublets are removed by circling the right area in the FSC-height vs area plot. Seeded single cells were then cultured using aforementioned medium or a modified medium with 20% FBS for two weeks. Culture medium was not changed until clear colonies were observed in some wells. If certain treatments are needed, i.e. double drug dose changes, they are initiated upon seeding the cells.

### Metaphase Analyses and Fluorescence in situ Hybridization (FISH)

Cells were blocked at metaphases by adding colcemid (KaryoMax, Thermo Fisher Scientific) at a final concentration of 0.05μg/ml followed by incubation at 37°C for 6-8 hours. Cells were then fixed using methanol:acetic acid (3:1). FISH slides were prepared by dropping fixed cells in a humid environment following the manufacture’s protocol provided for the BRAF/CCP7 FISH Probe Kit (Cytotest). Colored FISH images were taken and processed using confocal microscope Leica TCS SP8 X. Karyotype categorizations were based on the guidelines in Supplementary Fig. S1.

### qPCR BRAF Copy Number Assay

qPCRs for BRAF genomic DNA (gDNA) copy number measurement were performed by combining samples with PowerUp SYBR Green Master Mix (Applied Biosystems) in Optical 96-Well Reaction Plates (Applied Biosystems) with three technical replicates for each sample. Plates were then read by 7500 Real-Time PCR System (Applied Biosystems) using the standard cycling mode. Input templates for all samples were genomic DNAs extracted using DNeasy Blood & Tissue Kits (Qiagen). Unless specified, all qPCR runs used M249 parental as the reference sample and GAPDH as the endogenous control. All primers were ordered from Eurofins Scientific and their sequences are shown below.

BRAF Forward: 5’-TTTAGAACCTCACGCACCCC-3’ (intron 2) BRAF Reverse: 5’-TGTTGTAGTTGTGAGCCGCA-3’ (intron 2)

GAPDH Forward: 5’-CTGGCATTGCCCTCAACG-3’

GAPDH Reverse: 5’-AGAAGATGAAAAGAGTTGTCAGGGC-3’

### Comparative Genomic Hybridization (CGH) and Whole Genome Sequencing (WGS)

Genomic DNA of M249-P and M249-VSR cells were isolated by using DNeasy Blood & Tissue Kits (Qiagen). Samples were run on Agilent 6×80K array. The raw data was then processed by Cytogenomics software (Agilent Technologies). Nested genomic regions were flattened and .seg files were generated through R programming, followed by data visualization in IGV ^62^. Regions with large copy number changes were identified by comparing every segment in M249-VSR with the corresponding segment in M249-P. The same genomic DNAs were sent to PacGenomics for shallow WGS with coverage of 0.04. CNAs were inferred using Ginkgo ^63^, which contains a step that used bowtie ^64^ to align raw reads to hg19 genome.

### RNA-seq Analysis

Total RNA was isolated from M249-P and M249-VSR cells by using RNeasy Plus Mini Kit (Qiagen). Samples were sequenced on HiSeq3000 (Illumina) at 150bp paired-end. Raw data was then processed using Toil pipeline to output transcripts per million (TPM), including the STAR algorithm that aligned raw reads to GRCh38 genome ^65,66^. Following data trimming and log transformation, visualizations were done in R. For calculating allele frequencies of BRAF V600E, all RNA-seq fastq files were aligned using STAR, and the resultant bam files were processed according to GATK RNA-seq short variant discovery best practices until the step of haplotype calling ^67^. For visualization, we loaded base quality score recalibrated bam files to IGV.

### Immunoblotting and Antibodies

Cell lysates were prepared by using mRIPA buffer supplemented with PMSF, leupeptin and aprotinin. Western blots were performed using following antibodies: beta-actin (AC-15, Sigma-Aldrich), beta-actin (13E5, Cell Signaling Technology), BRAF (F-7, Santa Cruz Biotechnology), BRAF (C-19, Santa Cruz Biotechnology), Goat anti-Rabbit secondary antibodies (IRDye 680RD, LI-COR), Goat anti-Mouse secondary antibodies (IRDye 800CW, LI-COR). Images were directly output by Odyssey CLx Imaging System (LI-COR).

### Reverse Transcriptase (RT) -PCR and -qPCR

Total RNA was extracted from fresh cells using RNeasy Plus Mini Kit (Qiagen). Reverse transcriptions were then performed by using SuperScript VILO cDNA Synthesis Kit (Invitrogen). cDNA was then used for PCR and qPCR. Primers for detecting exon18-10 and exon9-10 junctions were the same as what previously published ^25^. The regular PCR was performed using Phusion High-Fidelity PCR Master Mix with HF Buffer (New England Biolabs). The PCR products that targeted exon18-10 and exon9-10 were then combined for each sample and run on 2% agarose gel. For qPCR, each sample was combined separately with PowerUp SYBR Green Master Mix (Applied Biosystems) and loaded on Optical 96-Well Reaction Plates (Applied Biosystems) in triplicate. Plates were then read by 7500 Real-Time PCR System (Applied Biosystems) using the standard cycling mode.

### Replica Plating Screen for DM-KDD Subpopulation

Each of 41 Single cells derived clones (SCs) of M249-VSR-HSR cells (cultured at 2μM VEM+SEL) was seeded in 6 wells of 96-well plates with the same cell number per well. Three wells of each clone were treated by 5μM VEM+SEL while the other three stayed at 2μM. After 6 days, cell viabilities were measured by CellTiter-Glo Luminescent Cell Viability Assay. 13 of 41 SCs were picked for a second round of the dose increase screen to confirm the findings. The viability of SCs was visualized by heatmaps using the R package ComplexHeatmap ^68^.

### Barcode-Based Clone Tracing

ClonTracer Barcoding Library ^43^ was purchased from Addgene. The plasmid pool was expanded by electroporation transformation. Lentivirus was made by transfecting 293T cells. M249-VSR-HSR cells were tested for their puromycin dose-response and multiplicity of infection curves. For the actual infection, 54 million M249 HSR cells were spin-infected in 12 well plate with 8μg/ml polybrene, followed by a six-day puromycin (0.3μg/ml) selection. Day 0 refers to the end of the selection. Next, cells underwent a standard culture growth period with kinase inhibitors present until the genomic DNA collection points on day 14 and day 35. The sequencing library was prepared by PCR amplification of barcode regions using the primer sequence provided by the manufacturer. The libraries were paired-end sequenced on Illumina NextSeq 500 at 75bp read length.

### Statistical Analysis

Significance of BRAF copy number qPCR is presented by t-distribution-based 95% confidence intervals. Doubling times for M249 SCs and bulk cells were calculated by fitting exponential growth curves, and their error bars were derived based on a previously published method ^69^.

## Acknowledgments

T.G.G. is supported by the NIH NCI P01 CA168585 and R01 CA222877, the Melanoma Research Alliance MRA (Team Science Award 691165), the UCLA SPORE in Prostate Cancer (NIH NCI P50 CA092131), the W.M. Keck Foundation, and the UCLA Eli and Edythe Broad Center of Regenerative Medicine and Stem Cell Research Hal Gaba Director’s Fund for Cancer Stem Cell Research. FISH microscopy was performed at the Advanced Light Microscopy/Spectroscopy Laboratory and the Leica Microsystems Center of Excellence at the California NanoSystems Institute at UCLA with funding support from NIH Shared Instrumentation Grant S10OD025017 and NSF Major Research Instrumentation grant CHE-0722519. Flow cytometry was performed in the UCLA Jonsson Comprehensive Cancer Center (JCCC) and Center for AIDS Research Flow Cytometry Core Facility that is supported by National Institutes of Health awards P30 CA016042 and 5P30 AI028697, and by the JCCC, the UCLA AIDS Institute, the David Geffen School of Medicine at UCLA, the UCLA Chancellor’s Office, and the UCLA Vice Chancellor’s Office of Research. Next generation sequencing was performed at Technology Center for Genomics & Bioinformatics (TCGB) at UCLA. CGH and low-pass WGS were performed at PacGenomics.

## Author Contributions

Conceptualization, K.S., and T.G.G.; Methodology, K.S., N.S., N.R., and T.G.G; Experiments, K.S., J.K.M., W.P.C., J.S., E.P., N.T.S., and N.R.; Data Analysis, K.S., and J.K.M.; Writing-Original Draft, K.S.; Writing-Review and Editing, K.S., J.K.M., K.P. and T.G.G.; Supervision and Project Administration, T.G.G.

## Notes

### Competing Interest Statement

T.G.G. reports receiving an honorarium from Amgen, having consulting and equity agreements with Auron Therapeutics, Boundless Bio, Coherus BioSciences, and Trethera Corporation. The lab of T.G.G. has a research agreement with BridgeBio Pharma, and has completed a research agreement with ImmunoActiva.

