## Supplemental Info for "Plasticity of extrachromosomal and intrachromosomal BRAF amplifications in mediating targeted therapy dosage challenges"

### Supplementary Information for "Plasticity of extrachromosomal and intrachromosomal BRAF amplifications in mediating targeted therapy dosage challenges"

Kai Song, Jenna K. Minami, William P. Crosson, Jesus Salazar, Eli Pazol, Niroshi T. Senaratne, Nagesh Rao, Kim Paraiso, and Thomas G. Graeber

**Contents**

**Supplementary Figures and Legends: S1-S10**

**Supplementary Table S1**

### Supplementary Figure S1

#### DM+ & HSR-

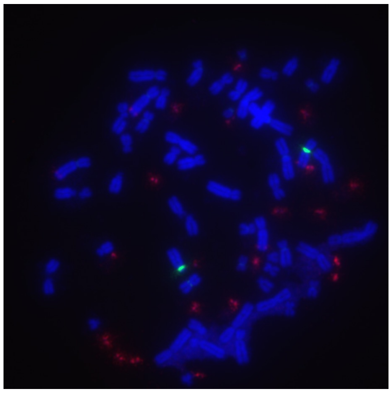

No. of DMs ~ 20

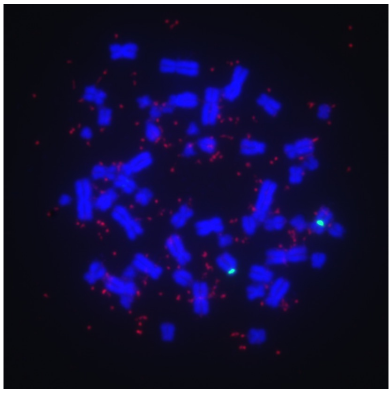

No. of DMs ~ 100

#### DM- & HSR+

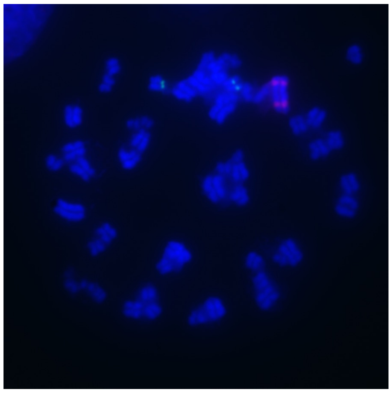

Two short HSRs on the same chromosome

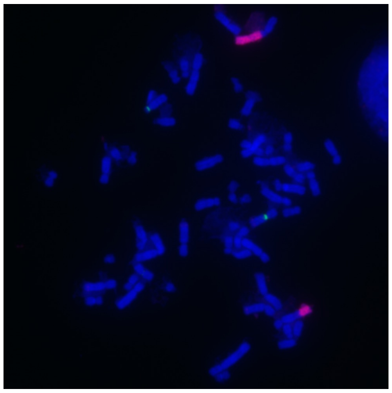

One long HSR and one short HSR on different chromosomes

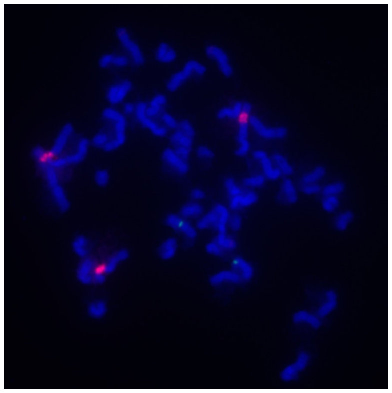

Three short HSRs on different chromosomes

#### DM+ & HSR+

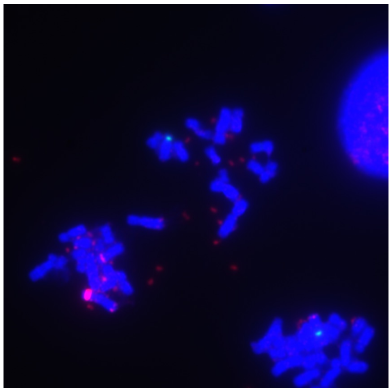

One short HSR and some DMs. HSRs are overexposed to make DMs bright

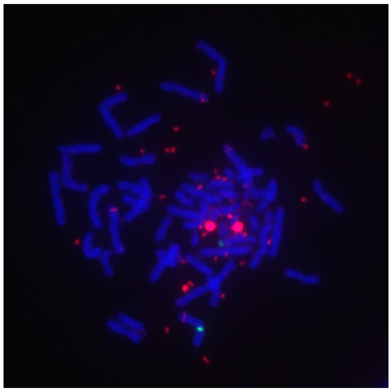

Two short HSRs and some DMs. HSRs are overexposed to make DMs bright

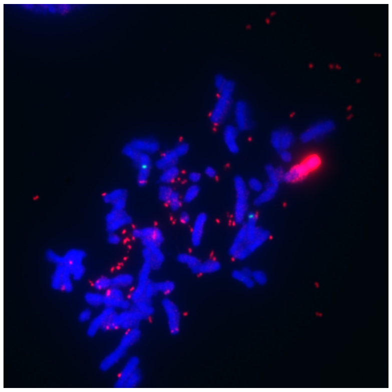

One long HSR and some DMs. HSRs are overexposed to make DMs bright

**Figure S1. BRAF FA karyotype categories and subcategories.** We divided all karyotypes into four primary categories: DM- & HSR-, DM+ & HSR-, DM- & HSR+, and DM+ & HSR+. Some categories have distinguishable sub-categories. Shown are representative FISH images of each BRAF FA category and sub-category. Red: BRAF. Green: centromere 7. Blue: DAPI.

### Supplementary Figure S2

M249-P

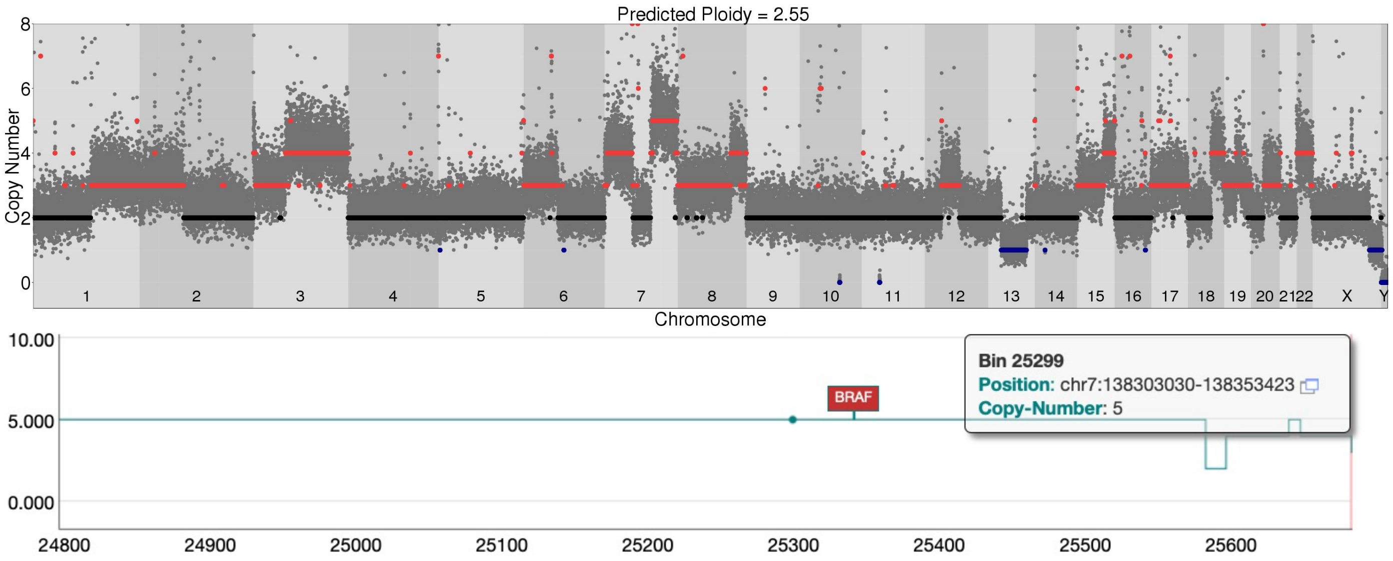


M249-VSR-DM

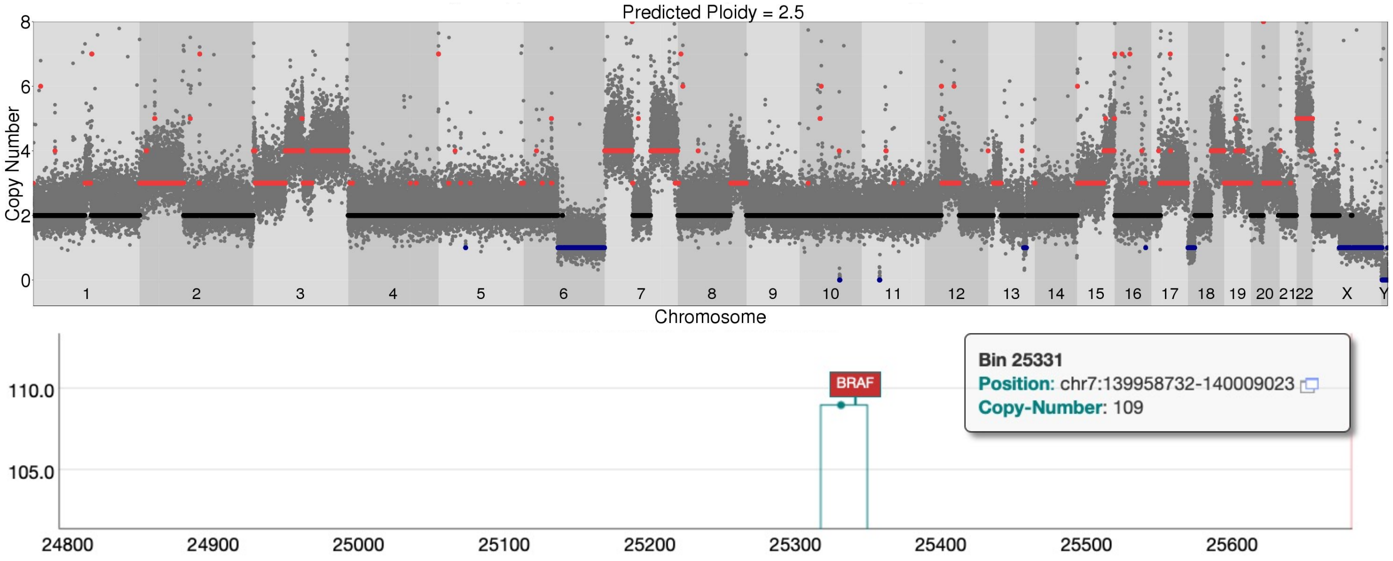


**Figure S2. BRAF DNA copy number amplification results confirmed by a second method.** Related to Fig 1. Whole genome sequencing (WGS)-based BRAF and genome-wide copy number results of M249-P and M249-VSR-DM cells were in line with comparative genome hybridization (CGH) results shown in Fig. 1E. Plotted is the whole genome CNA overview generated by Ginkgo. Below are the zoomed-in plots at the BRAF locus. Copy number values at the positions indicated by the green dots are shown in the inset boxes.

#### Supplementary Figure S3

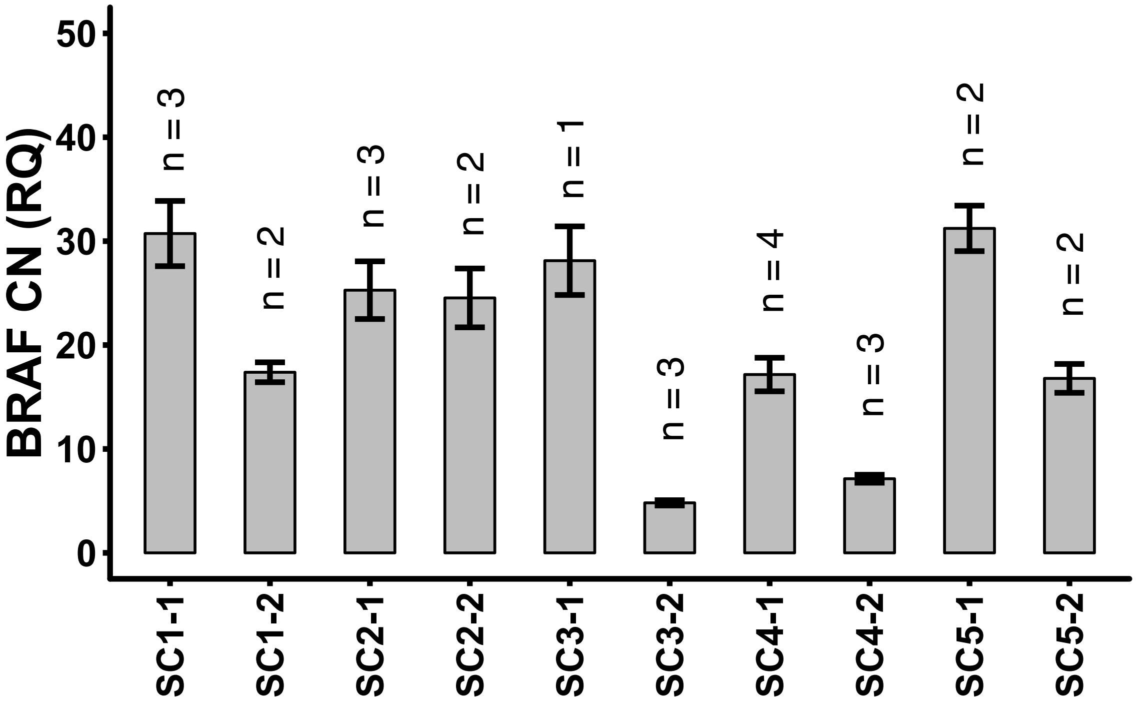


**Figure S3. Changes in BRAF copy number in single-cell-derived clones before and after three-month culture.** Related to Fig 2. M249-VSR cells were grown at the original 2μM dose of VEM+SEL. BRAF copy number was determined by qPCR. All values were normalized to GAPDH copy number of corresponding samples, and then to the values of the M249 parental cells. Plotted is the relative quantity (RQ) of BRAF copy number (CN) calculated by averaging multiple independent qPCR runs (n represents number of replicates). Error bars were calculated using propagation of errors.

### Supplementary Figure S4

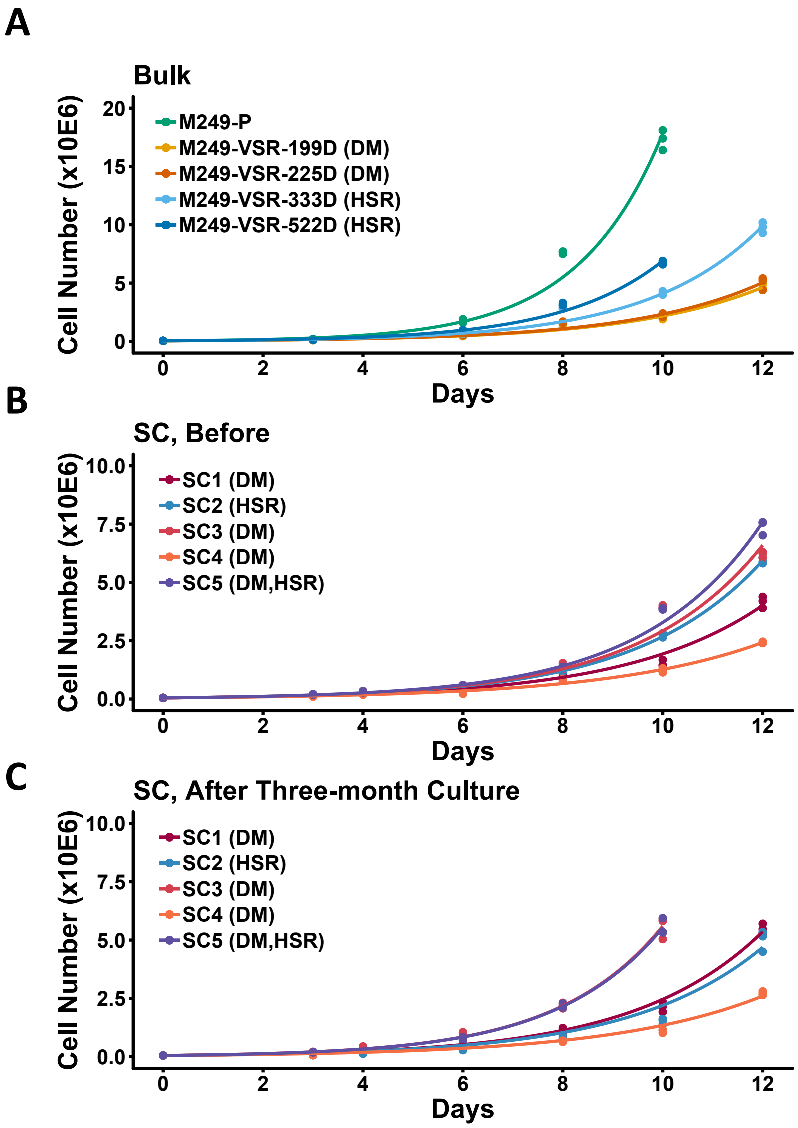


**Figure S4. Bulk MAPK inhibitor resistant melanoma cells displayed an increase in growth rate over time, while single-cell derived clones (SC) showed varying degrees of change in growth rate.** Related to Fig 2. Three technical replicates for each time point. **A**, Bulk M249-VSR cells. The days since establishment as a resistant subculture is indicated in the legend as xxxD. **B**, SCs shortly after single cell clone establishment. **C**, SCs after 3 months of culture. 0.05 million cells were plated in each well of 12-well plates, and cell numbers were monitored for a maximum of 12 days. Data points were fitted to the exponential growth curve $y = y_0 \cdot e^{kx}$ , where $y_0$ is the initial cell number, i.e. 0.05 million, y is the cell number at time x, and k is the rate constant.

### Supplementary Figure S5

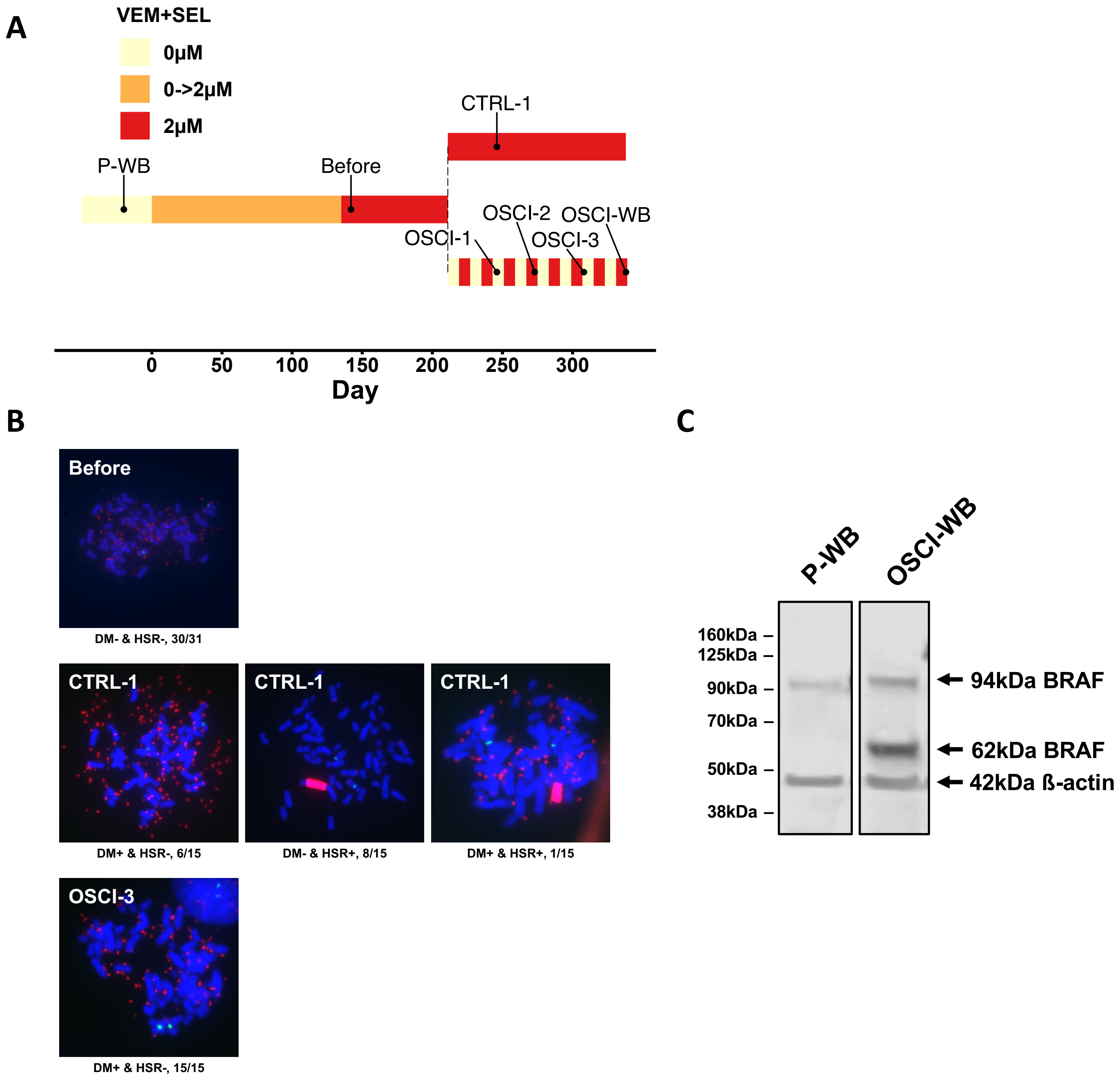


**Figure S5. Treating DM+ cells with oscillating doses of kinase inhibitors conferred a selection advantage for the DM+ & HSR- subpopulation.** Related to Fig 3. **A**, Oscillating and steady dose treatment schemes of M249-VSR-DM cells using VEM+SEL inhibitors. **B**, Representative FISH images for the sampling points indicated in (**A**). The fraction under each image represents the number of observations for this karyotype divided by total number of observations. Red: BRAF. Green: centromere 7. Blue: DAPI. **C**, Western blot results for M249 Parental sample and M249 VSR with oscillating dose (labeled in **A**).

### Supplementary Figure S6

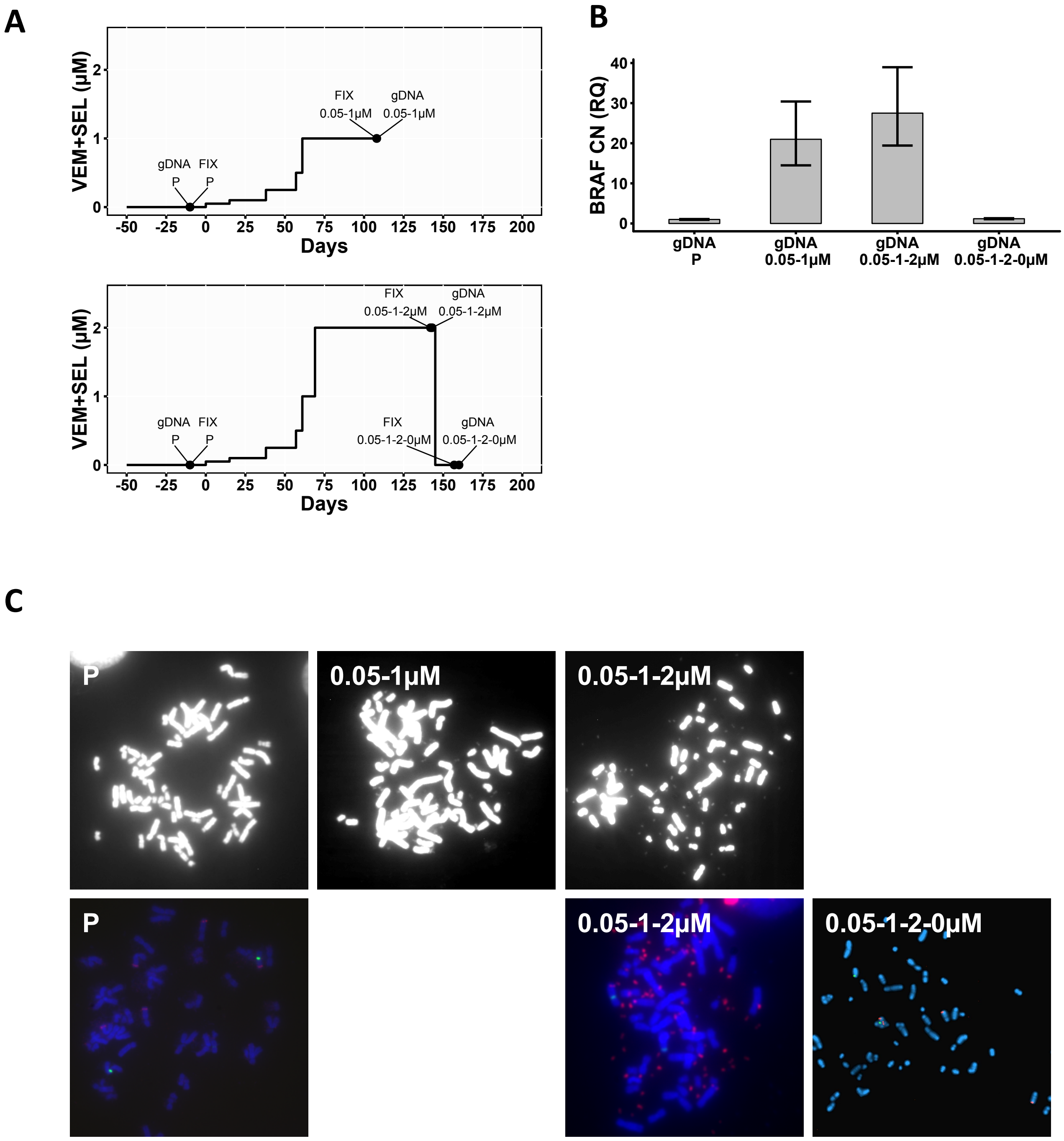


**Figure S6. Double drug withdrawal eliminated BRAF-carrying DMs in about 15 days.** Related to Fig 3. **A**, Treatment scheme of M249 parental cells with VEM+SEL. Points shown represent when cells were fixed and collected for genomic DNA (gDNA). Collections only occurred after cells were confirmed to have developed resistance against the current dose as determined by growth rates. **B**, qPCR results of relative BRAF copy number for the time points in (**A**). All values were normalized to GAPDH copy number of corresponding samples and then to M249 parental cells. Error bars represent t-distribution-based 95% confidence intervals from technical triplicates. CN: copy number. RQ: relative quantity. **C**, Representative metaphase spread images and FISH images for the time points in (**A**). Red: BRAF. Green: centromere 7. Blue: DAPI.

### Supplementary Figure S7

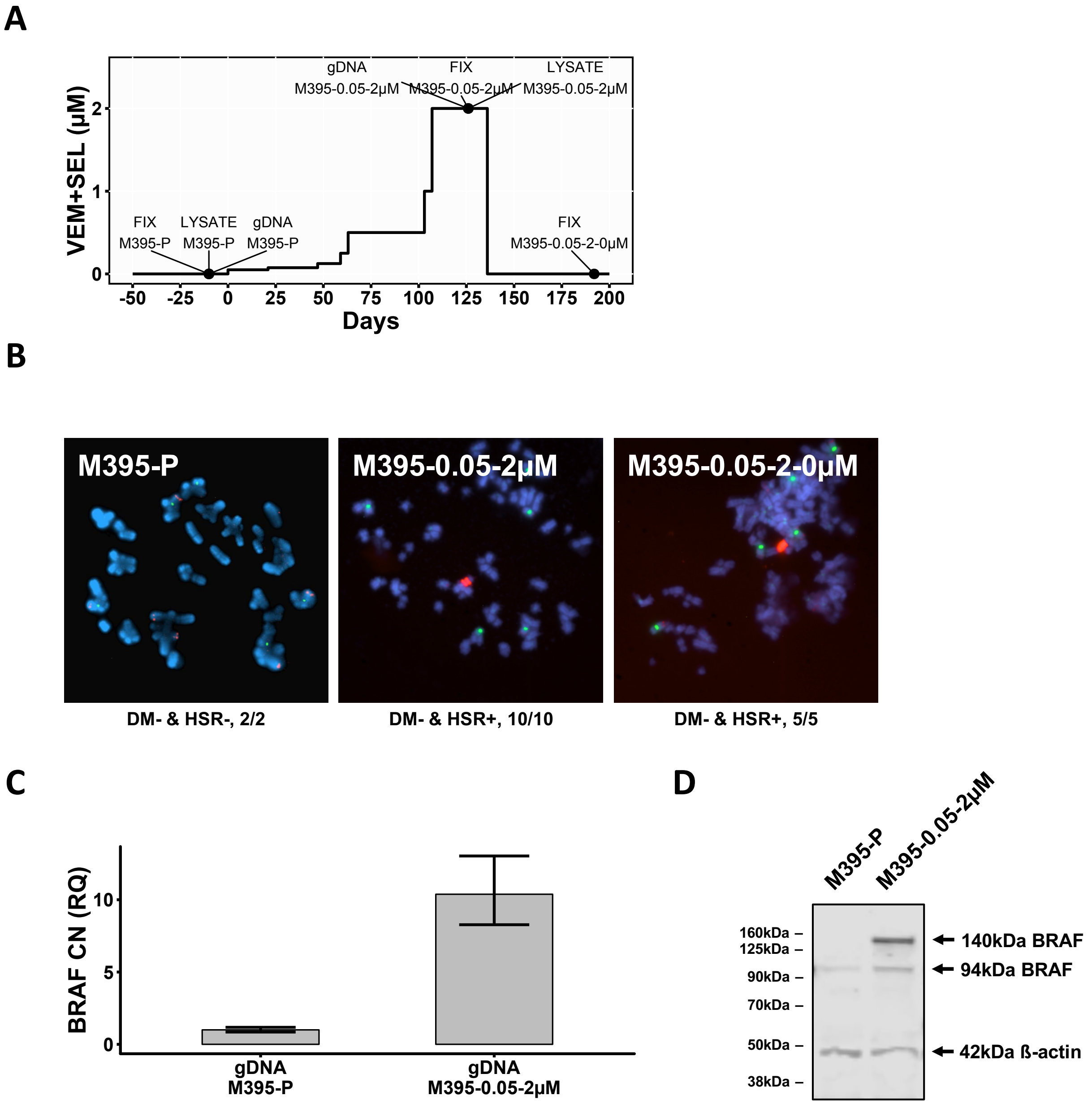


**Figure S7. Treatment of M395 melanoma cells with MAPK inhibitors led to BRAF amplification on HSRs co-occurring with BRAF kinase domain duplication.** Related to Fig 3. **A**, VEM+SEL treatment scheme starting from 0.05µM on M395-P (parental) cells. The points when cells were collected for gDNA (genomic DNA), fixation (FIX) and protein lysates (LYSATE) are labeled. **B**, Representative FISH images for all fixation time points in (**A**). The fraction under each image represents the number of observations for this karyotype divided by total number of observations. Red: BRAF. Green: centromere 7. Blue: DAPI. C, qPCR results of relative BRAF copy number in the samples for all gDNA collection points in (**A**). All values were normalized to GAPDH copy number of corresponding samples and then to parental cells. Error bars represent t-distribution-based 95% confidence intervals from technical triplicates. CN: copy number. RQ: relative quantity. D, western blot for certain samples in (**A**).

### Supplementary Figure S8

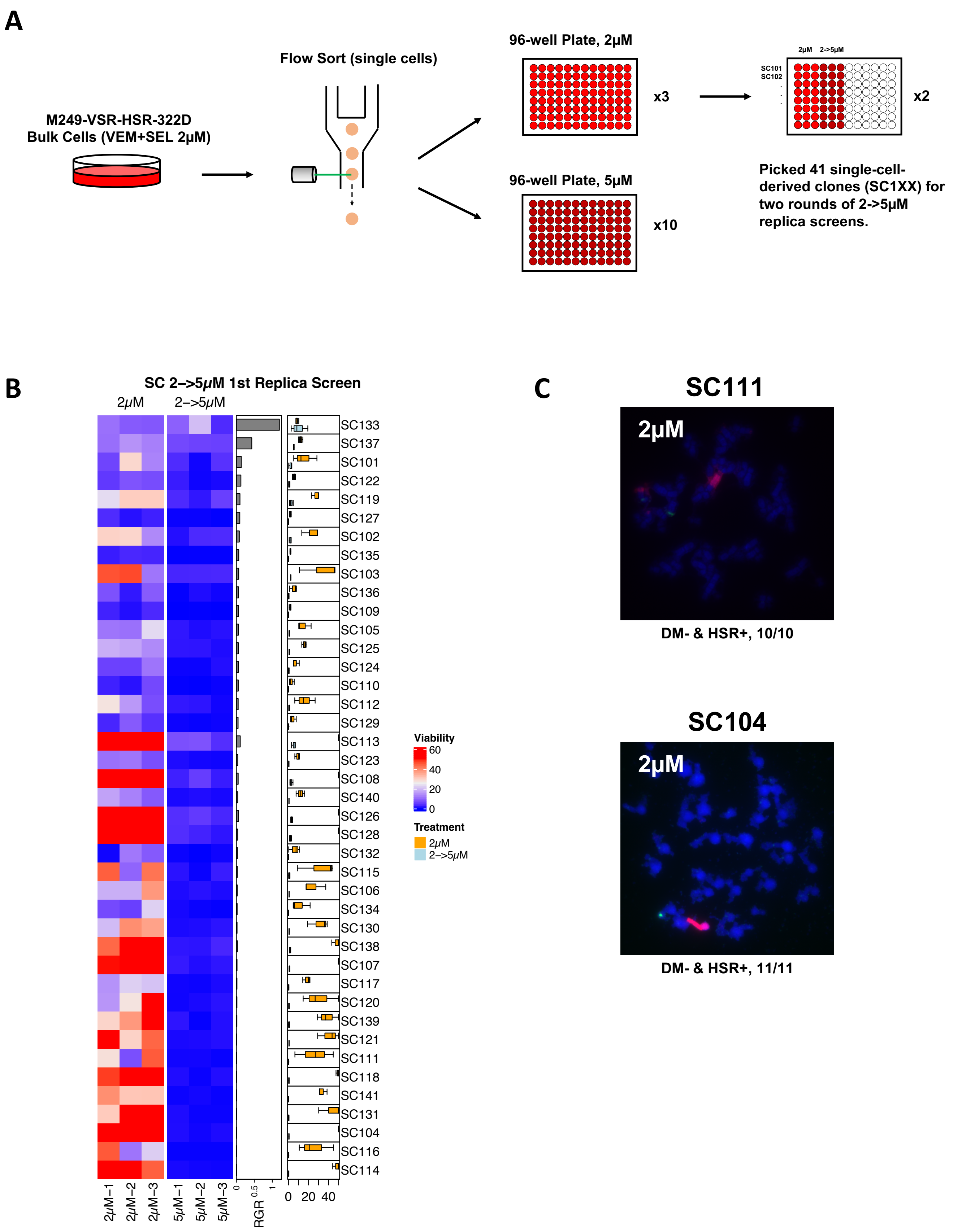

**Figure S8. Drug dose challenge characterization of single-cell-derived clones.** Related to Fig 4. **A**, Experimental design to generate single-cell-derived clones (SC1XXs) by sorting M249-VSR-HSR bulk cells on day 322, followed by two rounds of replica screens. **B**, As depicted in (**A**), acute 2 to 5μM VEM+SEL treatment on 41 SC1XXs was used to screen for clones that adapt to 5μM rapidly. The rows of the heatmap represent different SC1XXs ordered by relative growth rate (RGR), calculated by dividing the mean at 5μM by that at 2μM, in descending order. **C**, Representative FISH images of two SC1XXs at the lower tail of the heatmap in (**B**). The fraction under each image represents the number of observations for this karyotype divided by total number of observations. Red: BRAF. Green: centromere 7. Blue: DAPI.

### Supplementary Figure S9

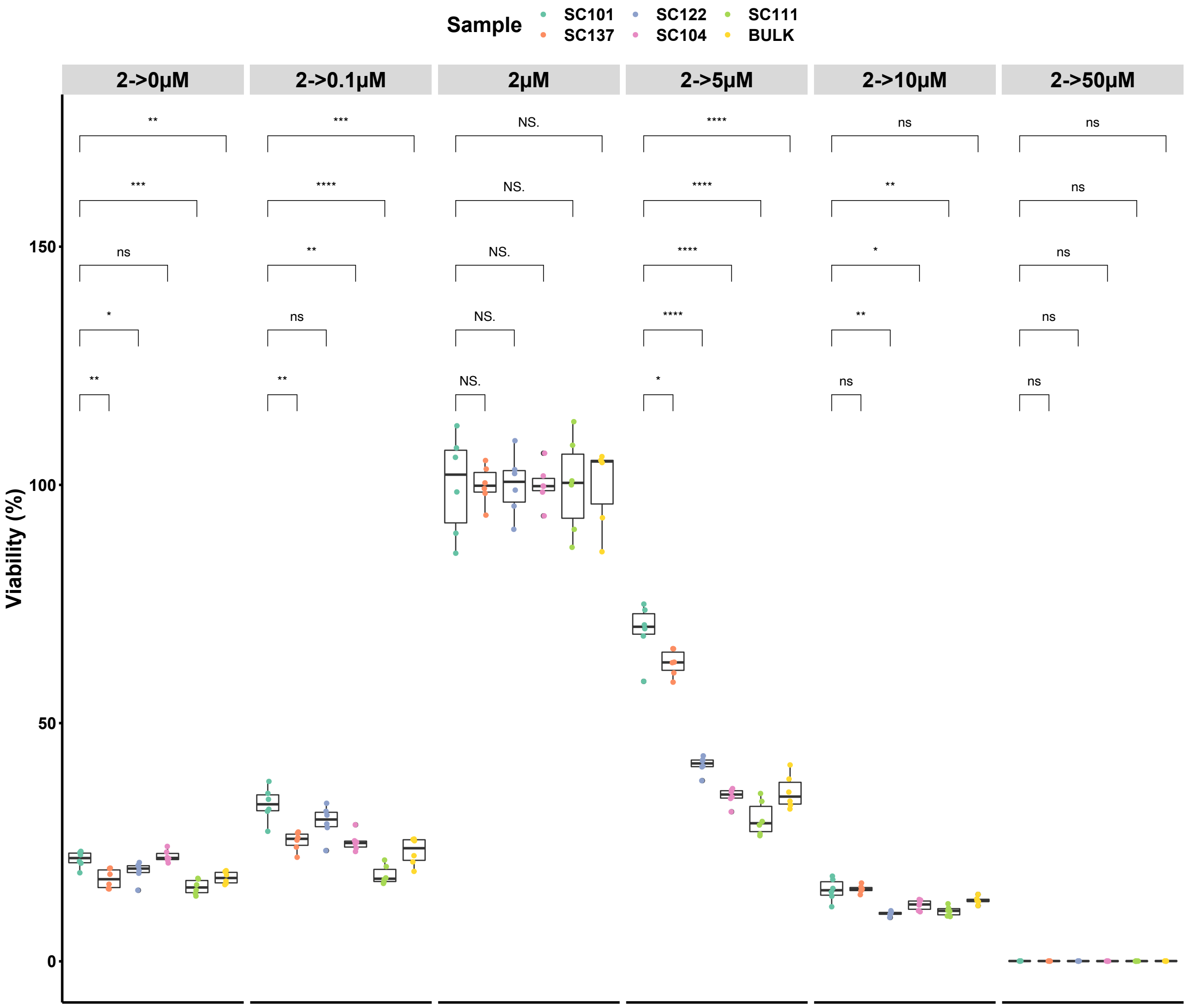


**Figure S9. The double-FA-mode (DM+ & KDD+) single-cell-derived clone SC101 tolerates a wider range of MAPK inhibitor dose challenges, compared to other SC1XXs.** Related to Fig 4. The indicated M249-VSR single cell clones and M249-VSR-HSR bulk cells initially cultured at 2μM of VEM+SEL were treated with various subsequent inhibitor doses for 4 days and then their viabilities were measured. All numbers are normalized to the corresponding viabilities at 2μM. Significance is calculated using the Student's t-test (n = 6).

#### Supplementary Figure S10

A

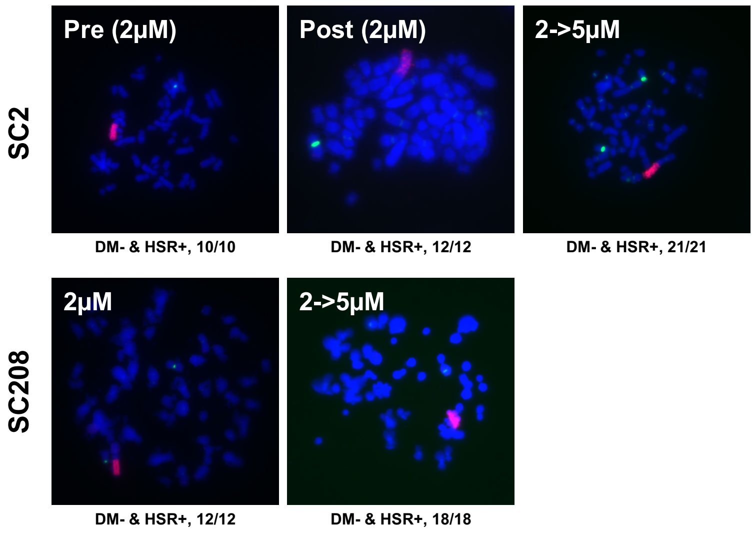


B

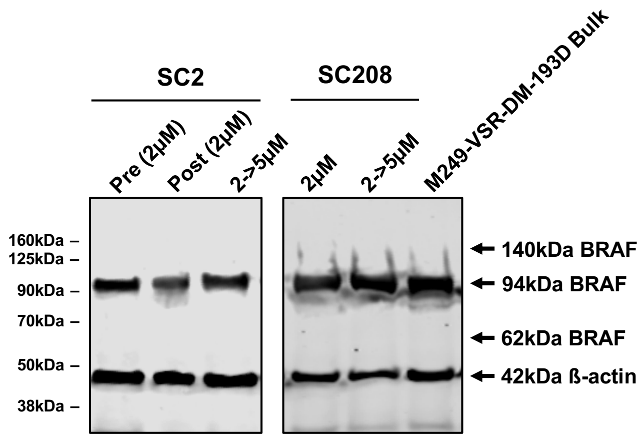


**Figure S10. MAPK inhibitor dose escalation applied to HSR-positive SCs did not result in the DM plus kinase domain duplication genomic configuration.** Related to Fig 4. **A,** representative FISH pictures of the DM- & HSR+ M249-VSR SCs, SC2 and SC208, dose escalated from 2µM to 5µM of VEM+SEL until they became resistant. The fraction under each image represents the number of observations for this karyotype divided by total number of observations. Red: BRAF. Green: centromere 7. Blue: DAPI. **B**, Immunoblot of BRAF samples in (**A**) showing no 140 kDa KDD band after the VEM+SEL dose increase.

**Supplementary Table S1. Karyotype evolution of single cell clones and the bulk cell population**

| | Karyotype circa day 250 | | | | Karyotype circa day 340 | | | | Summary | BRAF copy number change | Doubling time change |
|---|---|---|---|---|---|---|---|---|---|---|---|
| **DM** | - | + | - | + | - | + | - | + | | | |
| **HSR** | - | - | + | + | - | - | + | + | | | |
| **SC1** | 0 | 25 | 0 | 0 | 0 | 20 | 1 | 3 | DM+, some drift | 0.57x | 0.96x |
| **SC2** | 0 | 0 | 10 | 0 | 0 | 0 | 12 | 0 | HSR+, stable | 0.97x | 1.05x |
| **SC3** | 0 | 16 | 0 | 0 | 0 | 14 | 0 | 0 | DM+, stable | 0.17x | 0.86x |
| **SC4** | 0 | 21 | 0 | 0 | 0 | 13 | 2 | 3 | DM+, some drift | 0.42x | 0.98x |
| **SC5** | 0 | 0 | 0 | 16 | 0 | 0 | 13 | 2 | DM+ HSR+, transitioning to HSR+ | 0.54x | 0.89x |
| **Bulk** | 0 | 1 | 4 | 1 | 0 | 0 | 13 | 0 | mixed, transitioning to HSR+ | 0.91x* | 0.87x |

Summary of the changes in karyotype, BRAF copy number and doubling time for M249 VSR bulk and SCs over a three-month culture. Numbers under Karyotype columns are the number of karyotypes analyzed by FISH. The changes are calculated by dividing the latter date by the former. *The change of BRAF copy number for the bulk sample is calculated by dividing the value at circa day 4 by day 250.
